# Spatial fingerprinting: horizontal fusion of multi-dimensional bio-tracers as solution to global food provenance problems

**DOI:** 10.1101/2020.10.01.322453

**Authors:** Kevin Cazelles, Tyler Zemlak, Marie Gutgesell, Emelia Myles-Gonzalez, Robert Hanner, Kevin S. McCann

## Abstract

Building the capacity of efficiently determining the provenance of food products represents a crucial step towards the sustainability of the global food system. Whether it is for enforcing existing egislation or providing reliable information to consumers, technologies to verify geographical origin of food are being actively developed. Biological tracers (bio-tracers) such as DNA and stable isotopes have recently demonstrated their potential for determining provenance. Here we show that the data fusion of bio-tracers is a very powerful technique for geographical provenance discrimination. Based on 90 individuals of Sockeye salmon that originate from 3 different areas for which we measured 17 bio-tracers, we demonstrate that increasing the combined bio-tracers results in stronger the discriminatory power. The generality of our results are mathematically demonstrated under simplifying assumptions and numerically confirmed in our case study using three commonly used supervised learning techniques.

## 1. Introduction

> “*The future of food authentication and food quality assurance critically depends on combining chemometrics, computational analytical methods, and bioinformatics in processing and interpreting the data obtained through analytical technique*.^*1*^*”*

Today’s global food system is a collection of highly inter-connected trade networks that span a myriad of organizations and geographies.^2–5^ While such a system represents an important and perhaps necessary mechanism for meeting demands for nutritious and affordable food,^4,6^ it also represents a complex web of activity that carries with it a number of inherent challenges such as sustainability and transparency.^3,7^ Most food items travel thousands of kilometers^8^ changing form and ownership several times before reaching a consumer’s plate.^5^ Without proper labelling – such as Country of Origin Labelling (COOL) regulation – the consumer does not have the capability to accurately identify where their food originated and thus cannot make informed decisions about the products they are buying.^9^ Unfortunately, tracing food commodities back to their respective origin is a formidable task, which can only be tackled by a robust traceability system integrated along the entire food supply chain.^10^

The introduction of new technology (e.g. Wireless Sensor Network and Radio Frequency IDentification, blockchain) and ad-hoc recommendations (ISO 9000, Codex Alimentarius, etc.) represent indispensable tools to monitor and secure different food chain stages.^10,11^ However, there is one common limitation that is stopping us from realizing robust provenance-based value chains – the ability to verify traceability information.^1^ Consequently, a vigorous research effort has been geared towards the development of methods to authenticate type and origin of food commodities, such as sensory analysis and chromatographic techniques.^1,13,14^ One promising avenue is the use of bio-tracers, i.e. biological features (e.g. DNA, trace-elements, metabolomic compounds, stable isotopes, etc.) that vary with (and thus reflect) the environment an individual is living in, to create fingerprints that can recognize different food products. For instance, DNA barcodes have been used for over a decade to uncover fraudulent labelling practices in the seafood industry.^15–19^ Similarly, stable isotopes have shown a lot of promise for authenticating the origins of various food products including olive oils, cheese, meat and fish.^20,21^

In food product authentication, classes of bio-tracers are often employed independently of each other and “vertical” bio-tracer strategies (i.e., using different markers within a given class of bio-tracers) still prevail to adjust the granularity of information being sought. For example, small DNA sequence fragments of the mitochondrial cytochrome c oxidase I gene (COX-1) are enough to identify a fish fillet to species,^19^ but the genome coverage required is larger when trying to discriminate sub-populations where genetic variability is much smaller.^17^ Similarly, increasing the number of stable isotopes used has repeatedly been shown to be a powerful approach to determine the provenance (i.e. the origin) of numerous food products, even at fine spatial scales.^20,22–24^ Interestingly, despite the evidence of important gains in discriminatory power brought by vertical data fusion, the general reasons behind such success are rarely discussed. Furthermore, it remains unclear whether this gain extends beyond one class of bio-tracers, i.e. the potential of “horizontal” strategies for food authentication remains to be established.^1,25^

In what follows we discuss the efficiency of combining information from different bio-tracer (vertically and horizontally) for food authentication with a specific focus on provenance. Our goal here is threefold. First, we explain how to use multiple bio-tracers to create spatial fingerprints. Second, we show that increasing the number of bio-tracers for authentication increases authentication performance. Third, we provide a relatively simple explanation for why data fusion is always a winning strategy and comment on the potential of horizontal strategies. We support our arguments (which are mainly mathematical, see Supplementary Notes) by comparing how well a set of bio-tracers perform when trying to assign Sockeye Salmon (*Oncorhynchus nerka*) to 3 geographically distinct fisheries: British Columbia, Canada; Kamchatka Peninsula, Russia; and Alaska, United States (Fig. 1). The set of bio-tracers included 3 isotopes (*δ*^15^N, *δ*^13^C and *δ*^34^S) and 14 fatty acids for a total of 17 bio-tracers spanning two different classes. The Sockeye fishery itself presents an interesting model because sustainability practices vary somewhat geographically. The entire Alaskan fishery is certified by the Marine Stewardship Council (MSC) – the most rigorous and widely recognized eco-certification available. The Canadian fishery was also recognized by MSC as sustainable, until 2019 when the Canadian Pacific Sustainable Fisheries Society (CPSFS) decided to self-suspended its MSC certification for many salmon species, including Sockeye.^26^ While some fisheries in Russia received certification from the MSC, many remote fisheries in Eastern Russia are under threat due to extractive industries, loss of habitat and large-scale poaching.^27^ Much of this is thought to be driven by linkages to organized crime in east Asian markets.^28,29^ Therefore, building the capacity to distinguish high-level geographic origins of Sockeye is of particular relevance to the sustainability of Sockeye fishery and food provenance interests in general.

**Figure 1:**
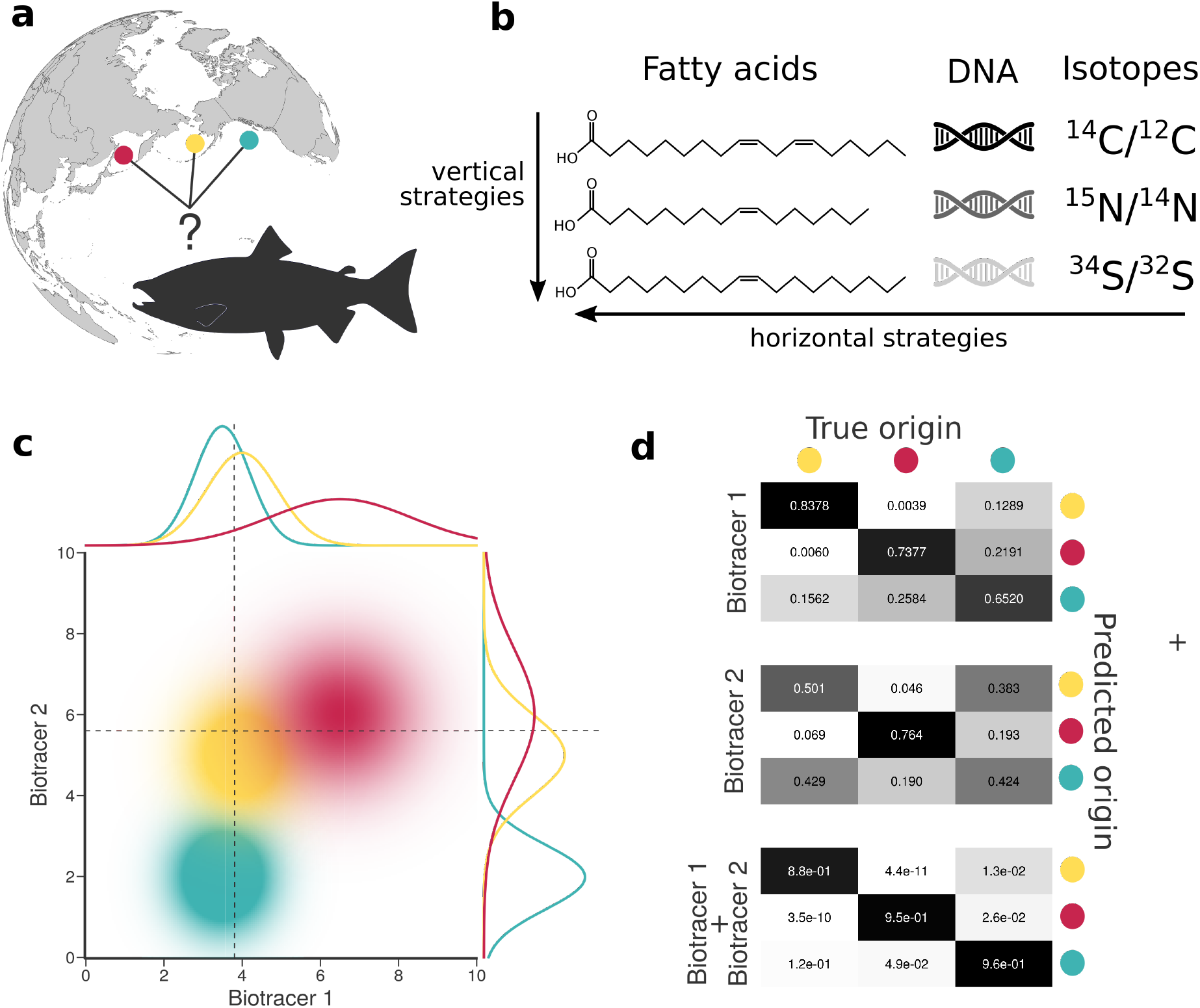
Combining bio-tracers to improve the determination of samples’ provenance. **a**, Sockeye salmon (*Oncorhynchus nerka*) samples of this study originate from three potential origins, namely Alaska, United States (yellow); British Columbia, Canada (cyan) and Kamchatka Peninsula, Russia (magenta). **b**, We examine the efficiency of horizontal strategies that combine several classes of bio-tracers as opposed to vertical strategies that focus on one specific class. **c**., While using a single bio-tracer to discriminate the true origin of a sample (distributions on top and right of the chart, dotted line depict bio-tracer values of a sample) may prove difficult, combining bio-tracers (colored areas) enhances the performance of the inference process. **d**, This is also shown with confusion matrices obtained using a classifier that uses only the first bio-tracers (top), only the second one (middle) or the combination of the two (bottom).

## 2 Results

### 2.1. Multiple bio-tracer distributions as spatial fingerprints

Bio-tracers developed over the past few decades uncover a variety of relationships between space and individuals. For instance, traces of local adaptations are encrypted in individual’s DNA.^30,31^ Stable isotope ratios and trace elements concentrations in tissues correlate with the surrounding physical environment such as soil and water.^25,32,33^ Similarly, the biotic component of the environment is reflected in the diet of an individual and thus in several biological indicators such as fatty acid profiles.^34^ Hence, the diversity of bio-tracers is a palette of authentication approaches^24,35^ focusing on complementary facets of the biology of a species to reveal its provenance. Therefore, taken together, all those pieces of information constitute spatially-distinct fingerprints with a high potential for food authentication.^1^

We created such spatial fingerprints of increasing complexity by combining up to 17 bio-tracers for our three regions of interest (see Fig. 1) and then evaluating their performances (on a different set of samples) to correctly determine the origin of a sample (see Methods). In order to cover a wide range of statistical approaches – reflecting current and emerging practices – we selected three classification approaches : Latent Discriminant Analysis (LDA), Naive Bayes Classifier (NBC) and Multi-Layer Perceptron (a class of artificial neural network ; hereafter MLP).

### 2.2. The more bio-tracers the better

For the three regions considered, increasing the number of bio-tracers always increased the probability of correctly assigning a sample to its true origin (Fig. 2). The three statistical approaches considered show similar behavior, qualitatively, with MLP having the best performance (Fig. 2c). All three methods consistently exceed 90% of correct assignment when 12 or more bio-tracers are combined for Canadian and Russian samples. The three approaches also perform significantly less efficiently for Alaskan samples, which are geographically closer to the two other regions (Fig. 1). Interestingly, the same order applied in the data space: the distance between Russia and Canada (based on the Euclidean distance between group centroids) is the longest (4.819 vs 4.145 for Canada-USA and 2.536 for Russia-USA).

**Figure 2:**
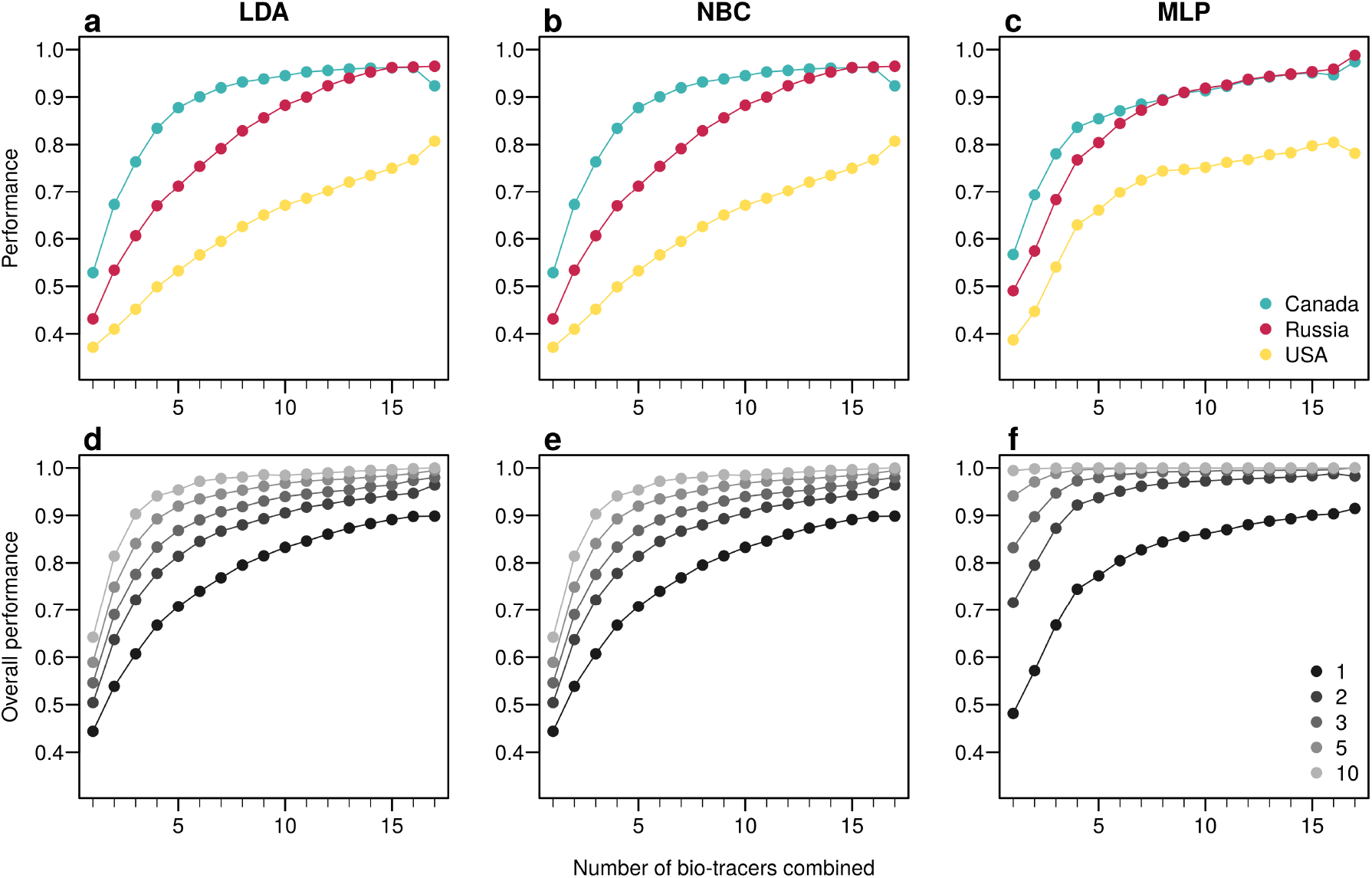
Increasing the number of bio-tracers considerably improves statistical performances. **a-c**, The probability of assigning one sample to its true origin increases as the number of bio-tracers employed increases for the three regions considered, namely Alaska (yellow), British Columbia (cyan) and Kamchatka Peninsula (magenta). **d-f**, The overall performance (i.e. the correct assigning any sample to its true origin) can also be improved by combining samples, assuming samples combined originate from the same region (e.g. individuals of the same lot). Points are colored according to the number of samples combined. These results are qualitatively similar for the three statistical approaches considered, which are Naive Bayesian classifier (NBC; **a, d**), Latent Discriminant Analysis (LDA; **b, e**) and a Multi-Layer Perceptron (MLP; **c, f**). In all panels, points represent performances averaged over up to 100,000 replicates (see Methods for further details).

The overall performance (i.e, the probability of correctly determining the provenance of a sample irrespective of its true origin), based on a single sample, from 1 to 17 bio-tracers increases from 0.444 to 0.898 for LDA, from 0.465 to 0.817 for NBC and from 0.482 to 0.915 for MLP (Fig. 2d-f). Moreover, performances are strongly improved when testing multiple individuals (Fig. 2d-f). It is worth noting that even in such case, employing more bio-tracers still provides more accurate predictions (Fig. 2d-f). Note that these results align fully with our analytical derivations (see Supplementary notes and Fig. S3). Also, increasing bio-tracers is very robust to noise addition, and this holds true for all three methods (Fig. 3). For instance, the overall performance of LDA with 5 bio-tracers and a very low level of noise added (10^−4^) is 0.710, but combining 15 bio-tracers with an addition of a noise with a level as high as 1 (a fairly strong noise addition) still yields better discriminatory power (0.758). Therefore, even if the measurement are known to be less accurate for some bio-tracers, they are likely worth being combined with others, assuming that the error is consistent among samples.

**Figure 3:**
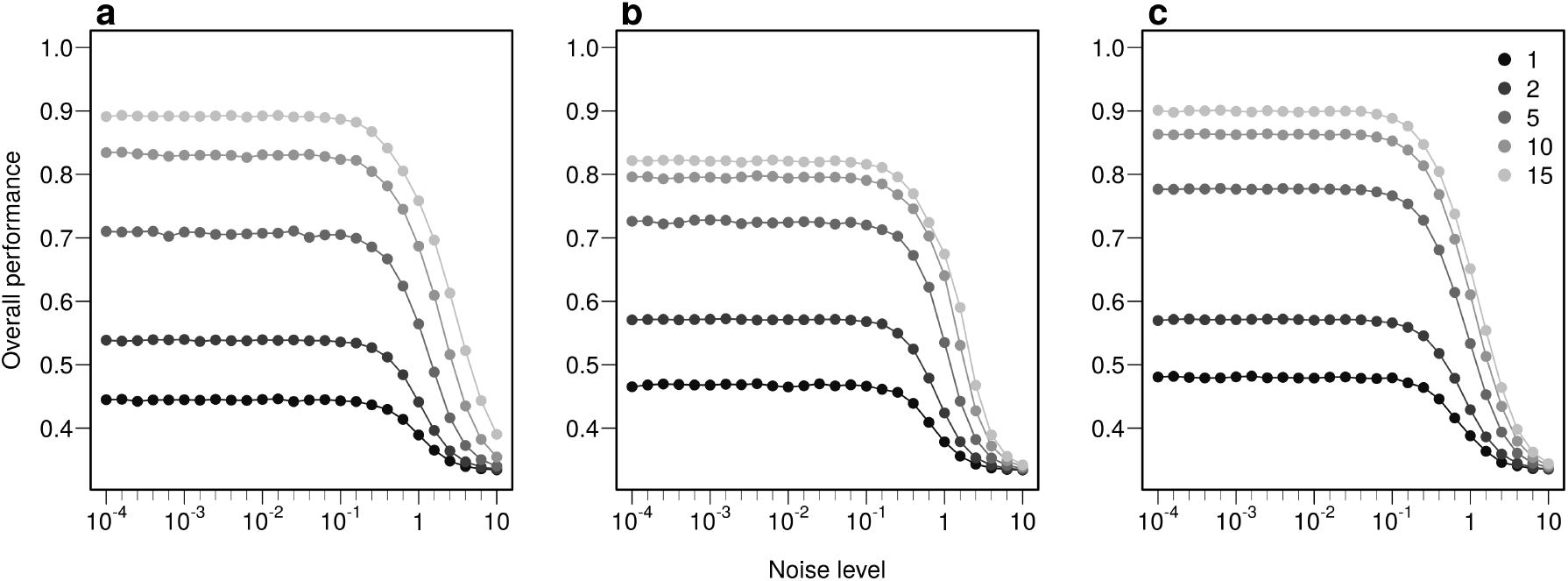
Combining bio-tracers is robust to noise addition. The probability of correctly determining the provenance of samples is evaluated for an increasing noise addition to the training data set. The lighter the gray, the more the number of bio-tracers combined. Note that prior to analysis, all bio-tracer values were scaled, thus a noise level of 1 represents a strong noise addition. The three panels correspond to three statistical approaches used: NBC (**a**), LDA (**b**), MLP (**c**).

Using the first axes provided by Principal Component Analysis (PCA) applied on the data set (see Methods) is a strategy that performs relatively well: across all three approaches, using up to the first 6 principal component axes is consistently better than the median of all the bio-tracer combinations we tested (Fig. 4). Also, as expected, the results obtained are similar when most or all bio-tracers are being used, except for NBC for which the PCA slightly negatively impacts the overall performance. Most importantly, for all methods, the axis order provided by a PCA (the first axis being the one that captures the most variance) does not necessary reflect their discriminate power. Hence, the three statistical methods show that the 5^th^ principal component axis provides a more important gain in performance than the 4^th^ one (Fig. 4). In general, combining only a few of the first principal component axes to authenticate food products, as frequently done,^36^ may be a sub-optimal approach as it can discard axes that carry less variance but more discriminatory power.

**Figure 4:**
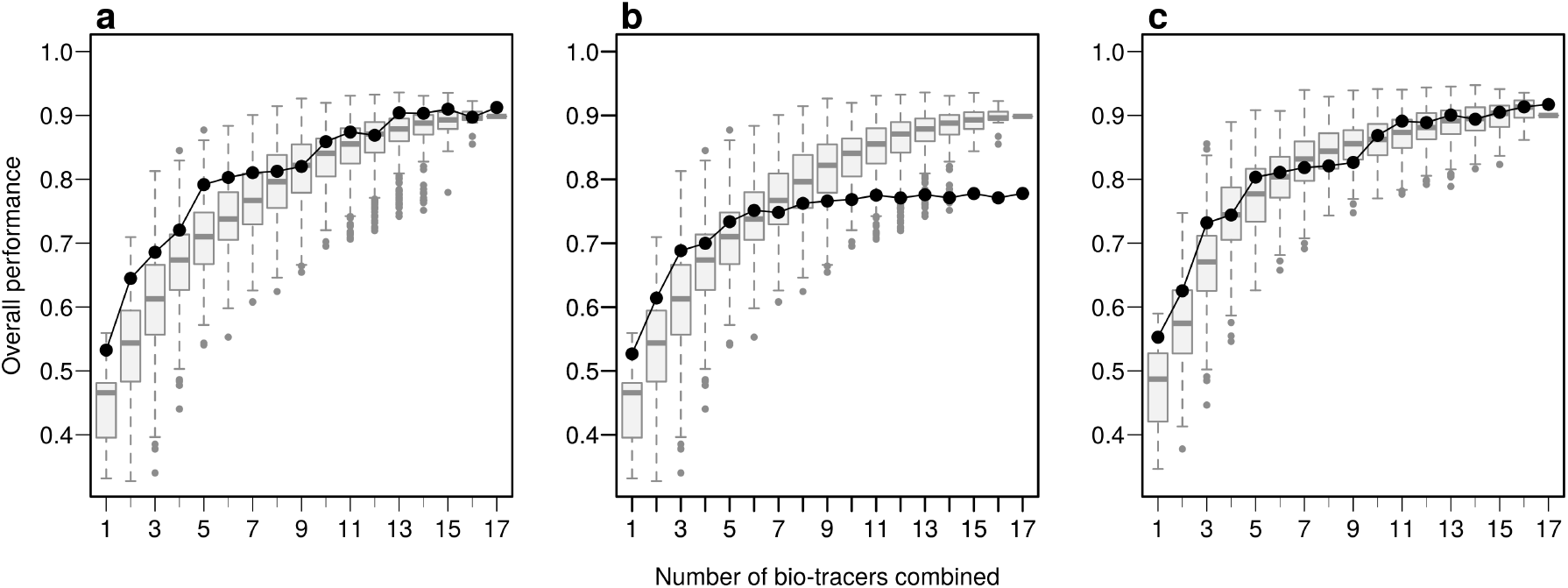
PCA does not necessarily provides axes with the maximum discriminatory power. Boxplots represent the probability of correctly determining the provenance of one sample for 500 combinations of bio-tracers (or all combinations if the total number of combinations is less than 500; see methods for details). Black lines and points represent results obtained when the first of principal component axes are being used. The three panels corresponds to three statistical approaches used: NBC (**a**), LDA (**b**), MLP (**c**).

### 2.3. An examination of the performances

Individually, the 17 bio-tracers have contrasting authentication performances (Fig. 5), this holds true for both classes of bio-tracers : *δ*^15^N and oleic acid (C18:1) alone perform well (0.547 and 0.548, respectively) whereas *δ*^13^C and linoleic acid (C18:2n-6) perform poorly (0.343 and 0.333). It is worth noting that the top 3 bio-tracers, based on individual performances, includes 2 fatty acids (oleic acid and docosapentaenoic acid, i.e. C22:5n-3) and one stable isotope (*δ*^15^N; see Fig. 5) and thus cover the two classes of bio-tracers. Note that even though we only show this for LDA (Fig. 5), this holds true for NBC (see Fig. S8) and MLP (see Fig. S11).

**Figure 5:**
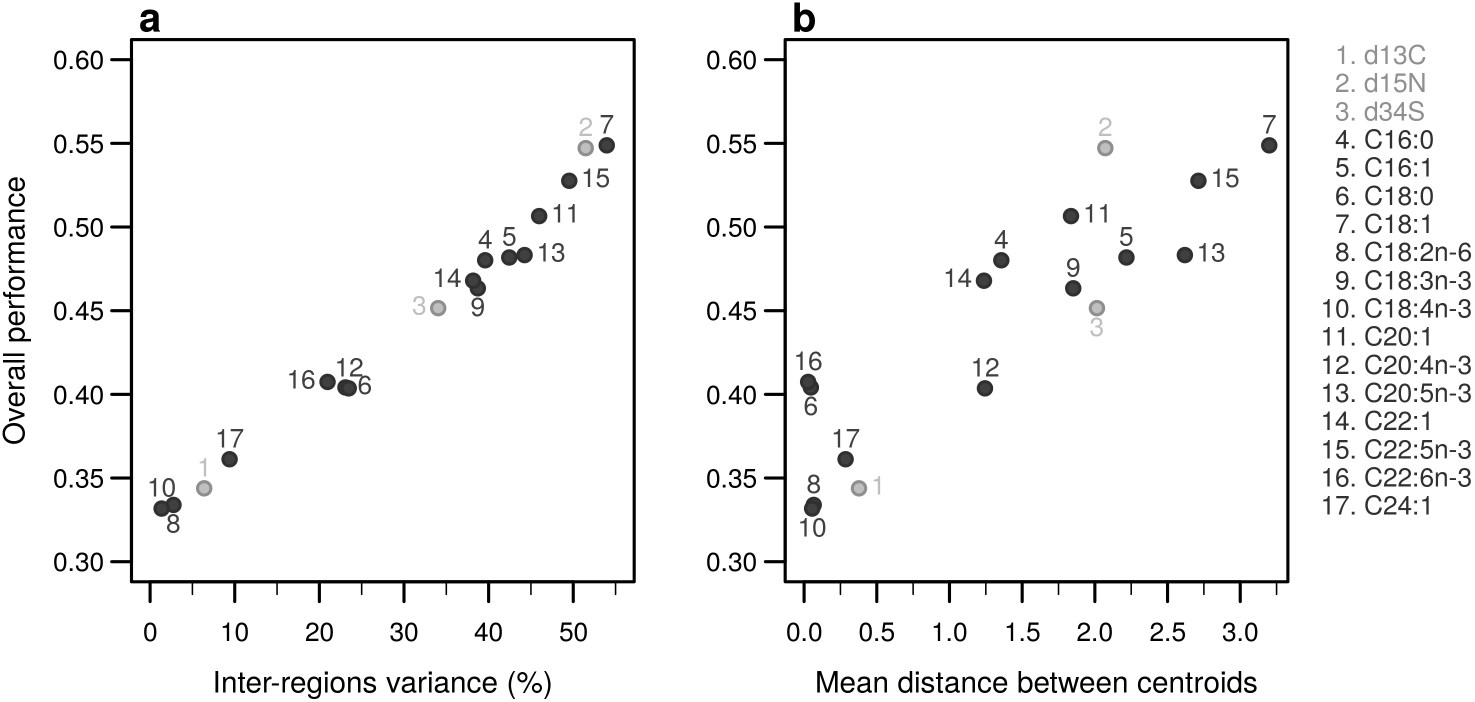
Performances of individual bio-tracers. Overall performances of all 17 bio-tracers (listed on the right) using LDA are plotted against the proportion of inter-regions variance (**a**) and the mean distance between all pairs of region centroids (**b**).

Interestingly, the overall performance of a pair of bio-tracers systematically outcompetes the best performing of the two bio-tracers included in the pair (see Fig. 6a for the results for LDA and Supplementary Fig. S9a for NBC and S12a for MLP). Similarly, the performance of combining the three bio-tracers is better than the best performing pair of bio-tracers that can be drawn from the triplet (see Fig. 6b and Supplementary Fig. S9b and S12b). Furthermore, the overall performance of a set of bio-tracers positively correlates with the performances of its subsets. Therefore using the best performing bio-tracers frequently yields a stronger discriminatory power (Fig. 6c-d and Supplementary Fig. S9c-d and S12c-d). This explains that the top 3 bio-tracers, and thus the two classes of bio-tracers, are systematically included in the best pairs and triplets (see Supplementary Tab. S1 and S2).

**Figure 6:**
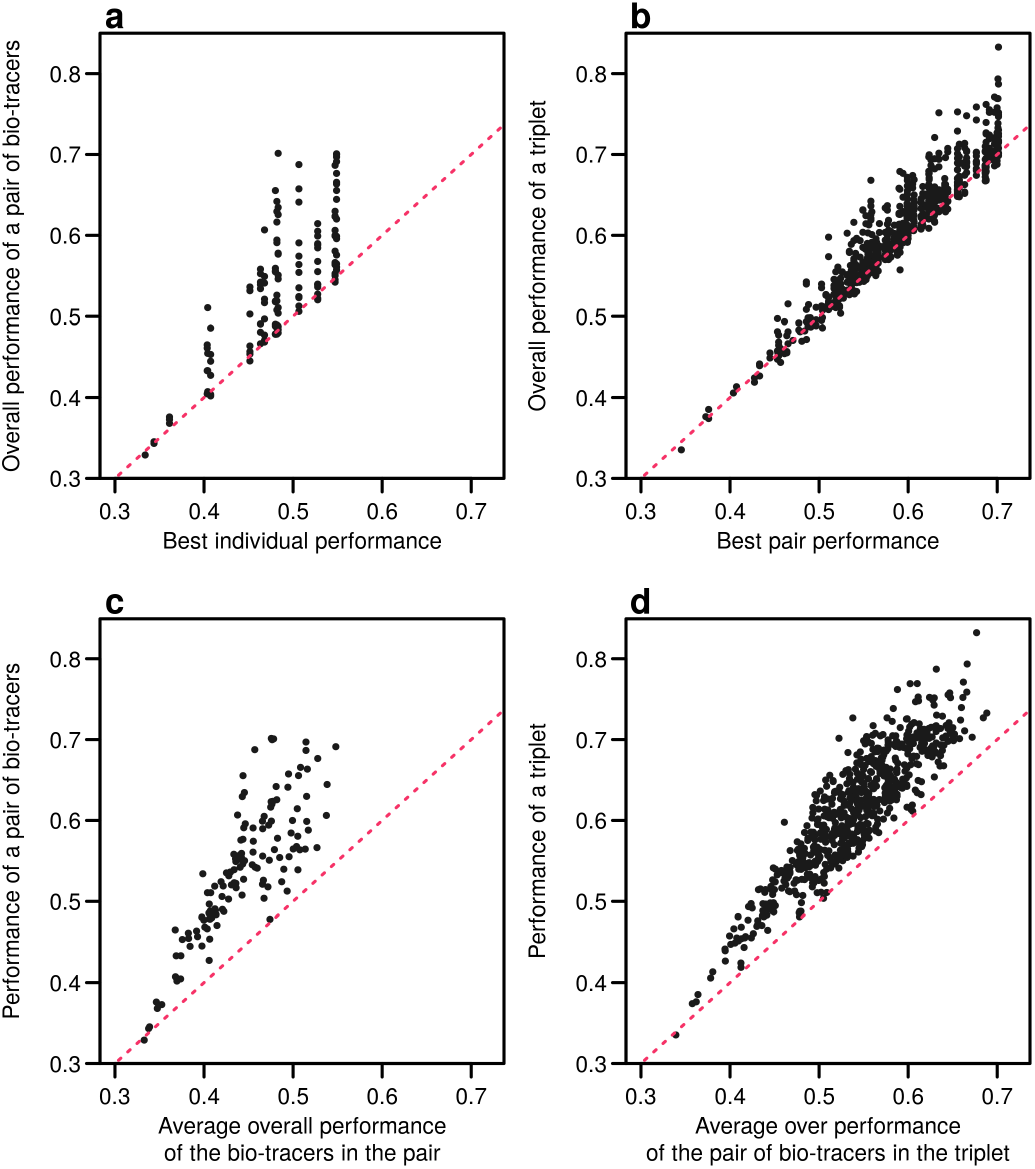
Including one more bio-tracers increases performance. Overall performances of all pairs of bio-tracers using LDA are plotted against the best individual performing bio-tracers of the pair (**a**) and their average overall performance (**c**). Similarly, overall performances of all triplets of bio-tracers are plotted against the best performing pair of bio-tracers of the triplet (**b**) and their average overall performance. Magenta dashed lines represent the 1:1 slope.

As expected, the percentage of inter-regions variance captured by a bio-tracer is strongly and positively correlated with its overall performance (Fig. 5a, Fig7-a,b). Even in 2 and 3 dimensions, simple non-linear least-squares regressions efficiently captures the variance of these relationship (R^2^=72.0% and 48.1% for LDA, respectively, see Fig. 7a-b and see Supplementary Fig. S10a-b and S13a-b for NBC and MLP, respectively). In one dimension, the mean Euclidean distance between region centroids efficiently summarizes one key geometrical results of the data space: the further apart the data points of different regions are, the stronger the discriminatory power (Fig. 5b). This result could be seen as a simple case of a more general one: the less overlap among regional hypervolumes (i.e. hypervolumes generated by data points of the different regions), the stronger the discriminatory power of a set of bio-tracers gets (Fig. 7c and Supplementary Fig. S10c and S13c). Notably, increasing dimensions is often an efficient way to reduce overlap among regions data points (see Fig. 7c, Supplementary Notes and Supplementary Fig. S10c and S13c).

**Figure 7:**
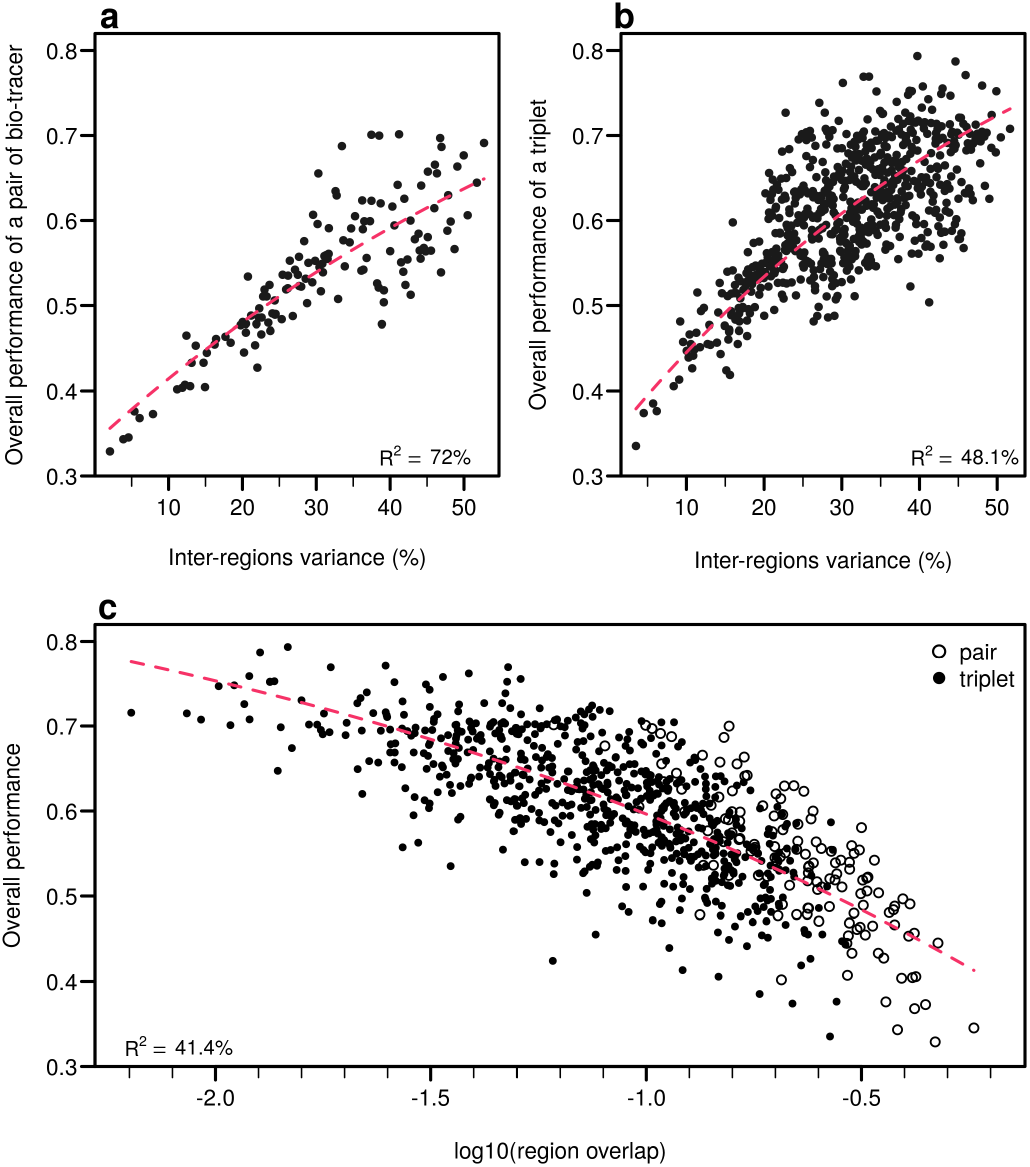
Efficient combinations of bio-tracers maximise inter-regional variance and minimize overlap of region data hypervolumes. **a-b**, For all combinations of 2 (**a**) and 3 (**b**) bio-tracers, the overall performances (with LDA) of sets of bio-tracers are plotted against their inter-regions variance. **c**, We present relationship between the proportion of overlap of data between regions and the overall performances for all pairs and triplets of bio-tracers. Magenta dashed lines represent the results of non-linear leas-squares regression and the corresponding R-squared are added at the bottom of every panel.

## 3. Discussion

Working in high dimensions for reliable authentication is already being used in our day-to-day life. For instance, face recognition algorithms use a high number of “abstract features” to recognize faces.^37,38^ Similarly, multi-messenger astronomy is experimenting with the fusion of electromagnetic radiation, gravitational waves, neutrinos and cosmic rays to observe and understand the universe through a new lens.^39^ Here we acknowledge similar potential for food authentication and clarify why data fusion can enhance the discriminatory power of traceability tools, and thus play a major role in food authentication in the foreseeable future, as other authors have already claimed.^1^ This is especially relevant for horizontal data fusion of bio-tracers as they are plentiful – some of which have just started to reveal their potential in tracing food products^40^ – and reflect various interactions between individuals and their immediate environment. Hence, together with technical advancements that trace movements of food products (such as blockchain), using bio-tracer based fingerprinting strategies to verify the origin of food product can contribute to making the food supply chain more transparent, more robust and eventually more sustainable.

Even though working in high dimension could be a very efficient approach, it also comes with its challenges : even if additional dimensions increase the discriminatory power of statistical classifiers, it comes at a cost as probability density estimates are more difficult and thus less accurate.^41^ This is where dimension reduction methods, such as PCA, can be utilized, as they allow for working in a simpler space with mathematically-desirable properties (e.g. uncorrelated axes concentrating the variance). However, one needs to bear in mind that what matters is to keep as much discriminatory power as possible and thus one should realize that, for instance, working with only the few first principal components may not always provide the best authentication tool as dimensions representing a low amount of total variance may still be of major importance to separate a pair of regions or more. Researchers should rather focus on statistical tools that reduce dimensions while maximizing discriminatory power, such as stepwise LDA.^33^ Fortunately, the recent boom in artificial intelligence research is bringing considerable methodological advancements in multivariate density estimation and dimension reduction.^42^

Taking advantage of data fusion can only be achieved if relentless efforts are made to acquire reliable data that would be securely archived (e.g. within a blockchain) while being widely accessible. This would require creating and maintaining ad hoc digital infrastructure. In our Sockeye example, we only needed 90 samples and 17 bio-tracers to cleanly differentiate a globally ubiquitous species by geographical region; however, we only covered 1 species across 3 spatially coarse regions – making high diversity and fine spatial scale applications will require more intensive data and probably the integration of additional classes of bio-tracers. While we did not consider DNA approaches beyond COI barcoding remains to be explored, we did explore DNA barcodes, well known for its species identification abilities,^43,44^ as a tool for spatial identification. Nonetheless, the genotypic variation at the COI gene was small and showed no spatial signal (see Fig. S.14). Moreover, here we did not investigate the temporal variations in bio-tracers distribution for the different regions which will be a critical step as this would determine the survey frequency required to maintain reliable spatial fingerprints. Overall, the sampling effort and the data required to extensively cover fishing areas experiencing food security concerns with numerous species of interest (and/or at risk) over long periods of time would certainly be bigger by several orders of magnitude.

Ultimately, standardizing sampling protocols, building large databases and employing powerful computational tools will allow researchers and national authorities to create dynamic maps of probability of origin for any food product to be tested.^32^ There are various strategies to improve food authentication, employing horizontal data fusion is clearly one of them. Fortunately, we are living in an era where major technical needs have been met, thus horizontal strategies can be employed immediately, but evidently their spread will depend on the balance between the cost of their application and the economical benefits for fishing industry, which vary across seafood products. That said, horizontal data fusion of bio-tracers could certainly be employed beyond the field of food authentication as it is a general principle where bio-tracers can be applied and combined to determine a wide array of biological properties, be it for determining the origin of a species or the structure of an entire food web.

## Acknowledgment

This project is part of the University of Guelph’s Canada First Research Excellence Fund project “Food from Thought”. This research was enabled in part by support provided by WestGrid (www.westgrid.ca) and Compute Canada (www.computecanada.ca). We would also like to acknowledge the generous support shown by many. We would like to specifically recognize Scott Wallace (David Suzuki Foundation), Liane Veitch (SeaChoice), and Kurtis Hayne (formerly with SeaChoice) for their inspiration, perspectives, and assistance coordinating samples. As well as Guy Dean (formerly with Albion Farms & Fisheries) for his foresight, passion for sustainability, and role in procuring samples. This research would not have been possible without you. Also, we are grateful to Marie-Hélène Brice for fruitful discussions throughout the writing of this paper.

## 4 Methods

### 4.1. Data

Muscle tissue trimmings were collected from 90 Sockeye salmon individuals from three different regions (30 individuals per region): British Columbia, Canada; Kamchatka Peninsula, Russia; and Alaska, United States were donated by Albion Farms & Fisheries Ltd. (now Intercity Packers Ltd.), Richmond, BC, Canada. All samples were derived from fillet trimmings to simulate a likely Quality Assurance/Quality Control scenario. Each muscle trimming was processed to obtain 2 muscle tissue samples for analyzing 17 bio-tracers of two classes: 3 stables isotopes (*δ*^15^N, *δ*^13^C and *δ*^34^S) and 14 fatty acids (listed in Fig. 3). One muscle sample from each fish was delivered frozen to the Lipid Analytical Services at the University of Guelph for fatty acid analysis using a combination of Bligh & Dwyer and Morrison & Smith methods.^46,47^ Individual FA weights (*µ*g/g) were converted to a % FA composition and fatty acids with >1% presence were retained as bio-tracers. The second muscle samples were dried at 70°C for 2 days and ground into a fine powder in preparation for stable isotope analysis. Tissue samples were sent to the University of Windsor GLIER Chemical Tracers Lab for isotopic analysis of *δ*^15^N, *δ*^13^C and *δ*^34^S (Windsor, ON, Canada). Importantly, all variables were centered and scaled before any statistical inference.

### 4.2. Statistical models

Among the large diversity of supervised-learning methods available, we chose three to reflect current and emerging practices in food authentication:

- Linear Discriminant Analysis^48^ (LDA), see 49 for a use case,
- Naive Bayesian Classifier^48^ (NBC), see 50 and 45 for examples,
- A Multi-Layer Perceptron^48^ (MLP), see 25 for a recent study.

For all three methods, we assessed the probabilities of correctly assigning a sample to its true origin (referred to as performance in main text) for every region (this corresponds to the diagonal of the confusion matrix) as well as the probability of assigning a sample to its true origin, irrespective of its true provenance (overall performance). Assuming that we have no prior expectation for the origin of a given sample, the overall performance corresponds to the mean of the diagonal of the confusion matrix.

### 4.3. Mathematical proof

In the Supplementary Notes, using Bayes’s rule, we demonstrated that increasing the number of bio-tracers combined almost surely increases the discriminatory power (performance) of a Naive Bayesian Classifier (NBC).

### 4.4. Simulations

For every simulation, we randomly selected 60 samples (20 per region) as training set and used the remaining samples (30 in total) to evaluate performances of combinations of bio-tracers (thus, the samples used to evaluate performances are different from the one use by the algorithm to create its own knowledge of the data). All simulations were replicated for all three selected classification approaches. We also evaluated the impact of respective size of the two data sets for and we show that gains of performances beyond 20 samples in the training set were marginal for all three methods (see Supplementary Fig. S8).

We evaluated the performances for an increasing number of bio-tracers (from single performances up to the combination of all the 17 bio-tracers available). For every number *p* of bio-tracers, we used 500 combinations of *p* bio-tracers. When there were less than 500 existing combinations, we used all of them. For every combination, we randomly chose 200 pairs of training and test sets, leading to up to 100,000 simulations for a given number of bio-tracers.

We also assessed the overall performances of the three approaches on the dataset ordered by a Principal Component Analysis (PCA). PCA is a statistical tool commonly used to reduce dimension,^41^ here PCA was used to transform our data set and obtained uncorrelated variables ordered according to the percentage of variance of the entire data set they capture.

To evaluate the robustness to noise, we added an increasing amount of white noise in the of the training set, i.e. for every simulation, we drew 60 values in a centered normal distribution of an increasing standard deviation (from 0.0001 to 10). For every simulation, we used 500 combinations of bio-tracers and 200 pairs of training and test sets (randomly chosen).

Finally, for all bio-tracers and all combinations of 2 and 3 bio-tracers, we computed the inter-regions variance as well as the distance between region centroids (coordinates of region centroids are the means of coordinates of all samples in a given region). We also computed the region data overlap. To do so, for the three regions studied, we computed the convex hull for all pairs and triplets of bio-tracers. Note that, in order to discard potential outliers, we only used 27 data points per region (90%), points included were the closest to their respective region centroid. We then computed the volume (or area) of all intersections between the three convex hulls, summed them and then divided the quantity thereby obtained by the total area (or volume) of the three convex hulls. Last, for all of these sets of bio-tracers we evaluated the performance bio-tracers using 1000 pairs of training and test sets (randomly chosen).

#### 4.4.1. Numerical implementation

For LDA, we used the R implementation available in MASS, function *lda()*.^51^ We implemented our own naive Bayesian classifier using R version 3.6.3^52^ and use *density()* for kernel density estimates.

Finally, we used the Julia library Flux.jl^53^ for the multi-layer perceptron (two dense layers and cross-entropy loss function). As this approach is data demanding, we used a simple data augmentation procedure : data in the training set were repeated and noise (random variables drew from a centered normal distribution of standard deviation *σ*) of various levels was added to it (as a centered normal distribution). After evaluating the performances under various augmentation scenarios (see Supplementary Fig. S6), we opted for 1000 repetitions of the data set and a noise level of *σ* = 0.01.

### 4.5. Code and data availability

Code and data will be released on GitHub upon acceptance.

## Notes

### Competing Interest Statement

The authors have declared no competing interest.

## References

1. Danezis, G. P., Tsagkaris, A. S., Brusic, V. & Georgiou, C. A. Food authentication: State of the art and prospects. Current Opinion in Food Science 10, 22–31 (2016).

2. Kneen, B. From land to mouth: Understanding the food system. (NC Press, 1993).

3. Godfray, H. C. J. et al. The future of the global food system. Philosophical Transactions of the Royal Society B: Biological Sciences 365, 2769–2777 (2010).

4. Ingram, J. A food systems approach to researching food security and its interactions with global environmental change. Food Security 3, 417–431 (2011).

5. Clapp, J. Distant agricultural landscapes. Sustainability Science 10, 305–316 (2015).

6. FAO. The future of food and agriculture Alternative pathways to 2050. (FOOD & AGRICULTURE ORG, 2018).

7. Béné, C. et al. When food systems meet sustainability Current narratives and implications for actions. World Development 113, 116–130 (2019).

8. Weber, C. L. & Matthews, H. S. Food-Miles and the Relative Climate Impacts of Food Choices in the United States. Environmental Science & Technology 42, 3508–3513 (2008).

9. Roebuck, K., Turlo, C. & Fuller, S. D. Canadians Eating in the Dark: A Report Card of International Seafood Labelling Requirements. (2017).

10. Aung, M. M. & Chang, Y. S. Traceability in a food supply chain: Safety and quality perspectives. Food Control 39, 172–184 (2014).

11. Badia-Melis, R., Mishra, P. & Ruiz-García, L. Food traceability: New trends and recent advances. A review. Food Control 57, 393–401 (2015).

12. Galvez, J. F., Mejuto, J. C. & Simal-Gandara, J. Future challenges on the use of blockchain for food traceability analysis. TrAC Trends in Analytical Chemistry 107, 222–232 (2018).

13. Luykx, D. M. A. M. & van Ruth, S. M. An overview of analytical methods for determining the geographical origin of food products. Food Chemistry 107, 897–911 (2008).

14. Danezis, G. P., Tsagkaris, A. S., Camin, F., Brusic, V. & Georgiou, C. A. Food authentication: Techniques, trends & emerging approaches. TrAC Trends in Analytical Chemistry 85, 123–132 (2016).

15. Wong, E. H.-K. & Hanner, R. H. DNA barcoding detects market substitution in North American seafood. Food Research International 41, 828–837 (2008).

16. Baker, C. S. A truer measure of the market: The molecular ecology of fisheries and wildlife trade. Molecular Ecology 17, 3985–3998 (2008).

17. Galimberti, A. et al. DNA barcoding as a new tool for food traceability. Food Research International 50, 55–63 (2013).

18. Shehata, H. R., Bourque, D., Steinke, D., Chen, S. & Hanner, R. Survey of mislabelling across finfish supply chain reveals mislabelling both outside and within Canada. Food Research International (2018) doi:10.1016/j.foodres.2018.12.047.

19. Shehata, H. R., Naaum, A. M., Garduño, R. A. & Hanner, R. DNA barcoding as a regulatory tool for seafood authentication in Canada. Food Control 92, 147–153 (2018).

20. Camin, F., Bontempo, L., Perini, M. & Piasentier, E. Stable Isotope Ratio Analysis for Assessing the Authenticity of Food of Animal Origin: Authenticity of animal origin food. Comprehensive Reviews in Food Science and Food Safety 15, 868–877 (2016).

21. Bontempo, L. et al. Characterisation and attempted differentiation of European and extra-European olive oils using stable isotope ratio analysis. Food Chemistry 276, 782–789 (2019).

22. Shin, W.-J. et al. Discrimination of the geographic origin of pork using multi-isotopes and statistical analysis. Rapid Communications in Mass Spectrometry 32, 1843–1850 (2018).

23. Chung, I.-M. et al. Geographic authentication of Asian rice (Oryza sativa L.) Using multi-elemental and stable isotopic data combined with multivariate analysis. Food Chemistry 240, 840–849 (2018).

24. Zhao, Y. et al. Authentication of organic pork and identification of geographical origins of pork in four regions of China by combined analysis of stable isotopes and multi-elements. Meat Science 165, 108129 (2020).

25. Wu, H. et al. Verification of imported red wine origin into China using multi isotope and elemental analyses. Food Chemistry 301, 125137 (2019).

26. Fiorillo, J. Canadian wild salmon fisheries quitting MSC program. IntraFish.

27. Centre, T. W. S. A review of IUU salmon fishing and potential conservation strategies in the Russian Far East. (2009).

28. Clarke, S. Trading tails: Russian Salmon fisheries and East Asian markets. TRAFFIC East Asia. (2007).

29. Clarke, S. C., McAllister, M. K. & Kirkpatrick, R. C. Estimating legal and illegal catches of Russian sockeye salmon from trade and market data. ICES Journal of Marine Science 66, 532–545 (2009).

30. Leys, M. et al. Spatial genetic structure in Beta Vulgaris subsp. Maritima and Beta Macrocarpa reveals the effect of contrasting mating system, influence of marine currents, and footprints of postglacial recolonization routes. Ecology and Evolution 4, 1828–1852 (2014).

31. Blower, D., Pandolfi, J., Bruce, B., Gomez-Cabrera, M. & Ovenden, J. Population genetics of Australian white sharks reveals fine-scale spatial structure, transoceanic dispersal events and low effective population sizes. Marine Ecology Progress Series 455, 229–244 (2012).

32. Ehleringer, J. R. et al. Hydrogen and oxygen isotope ratios in human hair are related to geography. Proceedings of the National Academy of Sciences 105, 2788–2793 (2008).

33. Zhao, H., Zhang, S. & Zhang, Z. Relationship between multi-element composition in tea leaves and in provenance soils for geographical traceability. Food Control 76, 82–87 (2017).

34. Happel, A., Rinchard, J. & Czesny, S. Variability in sea lamprey fatty acid profiles indicates a range of host species utilization in Lake Michigan. Journal of Great Lakes Research 43, 182–188 (2017).

35. Ishida, Y., Georgiadis, N. J., Hondo, T. & Roca, A. L. Triangulating the provenance of African elephants using mitochondrial DNA. Evolutionary Applications 6, 253–265 (2013).

36. Naccarato, A., Furia, E., Sindona, G. & Tagarelli, A. Multivariate class modeling techniques applied to multielement analysis for the verification of the geographical origin of chili pepper. Food Chemistry 206, 217–222 (2016).

37. Chen, D., Cao, X., Wen, F. & Sun, J. Blessing of Dimensionality: High-Dimensional Feature and Its Efficient Compression for Face Verification. in 2013 IEEE Conference on Computer Vision and Pattern Recognition 3025–3032 (IEEE, 2013). doi:10.1109/CVPR.2013.389.

38. Taigman, Y., Yang, M., Ranzato, M. & Wolf, L. DeepFace: Closing the Gap to Human-Level Performance in Face Verification. in 2014 IEEE Conference on Computer Vision and Pattern Recognition 1701–1708 (IEEE, 2014). doi:10.1109/CVPR.2014.220.

39. Bartos, I., Kowalski, M. & Institute of Physics (Gran Bretanya). Multimessenger astronomy. (2017).

40. Pimentel, T., Marcelino, J., Ricardo, F., Soares, A. M. V. M. & Calado, R. Bacterial communities 16S rDNA fingerprinting as a potential tracing tool for cultured seabass Dicentrarchus labrax. Scientific Reports 7, 11862 (2017).

41. Scott, D. W. Multivariate density estimation: Theory, practice, and visualization. (Wiley, 2014).

42. Liu, G., Liu, Q. & Li, P. Blessing of Dimensionality: Recovering Mixture Data via Dictionary Pursuit. IEEE Transactions on Pattern Analysis and Machine Intelligence 39, 47–60 (2017).

43. Hebert, P. D. N., Cywinska, A., Ball, S. L. & deWaard, J. R. Biological identifications through DNA barcodes. Proceedings of the Royal Society of London. Series B: Biological Sciences 270, 313–321 (2003).

44. Zemlak, T. S., Ward, R. D., Connell, A. D., Holmes, B. H. & Hebert, P. D. N. DNA barcoding reveals overlooked marine fishes. Molecular Ecology Resources 9, 237–242 (2009).

45. Bataille, C. P. & Bowen, G. J. Mapping 87Sr/86Sr variations in bedrock and water for large scale provenance studies. Chemical Geology 304-305, 39–52 (2012).

46. Bligh, E. G. & Dyer, W. J. A RAPID METHOD OF TOTAL LIPID EXTRACTION AND PURIFICATION. Canadian Journal of Biochemistry and Physiology 37, 911–917 (1959).

47. Morrison, W. R. & Smith, M. Preparation of fatty acid methyl esters and dimethylacetals from lipids with boron fluoride-methanol. 9.

48. Hastie, T., Tibshirani, R. & Friedman, J. The Elements of Statistical Learning. (Springer New York, 2009). doi:10.1007/978-0-387-84858-7.

49. Sun, S., Guo, B. & Wei, Y. Origin assignment by multi-element stable isotopes of lamb tissues. Food Chemistry 213, 675–681 (2016).

50. Wunder, M. B. Using Isoscapes to Model Probability Surfaces for Determining Geographic Origins. in Isoscapes (eds. West, J. B., Bowen, G. J., Dawson, T. E. & Tu, K. P.) 251–270 (Springer Netherlands, 2010). doi:10.1007/978-90-481-3354-3_12.

51. Venables, W. N., Ripley, B. D. & Venables, W. N. Modern applied statistics with S. (Springer, 2002).

52. R Core Team. R: A Language and Environment for Statistical Computing. (R Foundation for Statistical Computing, 2020).

53. Innes, M. Flux: Elegant machine learning with Julia. Journal of Open Source Software 3, 602 (2018).

