## Supplementary Information for "Spatial fingerprinting: horizontal fusion of multi-dimensional bio-tracers as solution to global food provenance problems"

### Supplementary Notes

**Conclusions** **12**

**References** **13**

In this first part of the Supplementary Information, we provide a mathematical proof of our main result, i.e. increasing the number of bio-tracers combined increase the probability of assigning a sample to its true origin. To do so, we use several strong simplifying assumptions which are necessary for the rigor of the demonstration. Importantly, numerical results of the main text fully align with the mathematical ones obtained below.

#### Objectives

We consider a sample of $n$ individuals that belong to one of $p$ existing populations and we aim at determining the population the sample originate from. In order to do so, $q$ properties (e.g. fatty acid profile) are measured for every individuals and we assume value distributions are known for every populations. For the sake of clarity, we refer to these measurements as bio-tracers and we assume that inferring the population of origin equates inferring the geographic origins of the sample. Below, we use a Bayesian framework to infer the geographic origin of a sample based on bio-tracers and to discuss how the properties of the value distribution affect the inference.

#### Notations and definitions

In what follows:

- $n$, $p$ and $q$ are three natural numbers other than zero;
- $\mathbb{N}_n$ is the set of natural numbers ranging from 0 to $n$, where $n \in \mathbb{N}^*$;
- $[X]$ denotes the probability of the event $X$ ($X$ being an event, or a realization of a random variable), and $[X|Y]$ probability of $X$ given $Y$;
- $\mathcal{N}(\mu, \sigma)$ denotes a Gaussian distribution of parameter $(\mu, \sigma)$;
- $\mathbb{E}(X)$ denotes the expected value of the random variable $X$.

We consider $q$ bio-tracers and $p$ populations. For any population $j$, $f_j$ denotes the probability density distribution (pdf) of bio-tracers values as follows:

$$f_j : D_1 \times ... \times D_q \to [0, 1]$$
$$\mathbf{x} \to f_j(\mathbf{x})$$

where $D_l$ denotes the support set of the $l^{th}$ bio-tracer (we use the same support set for all areas). Now let $\mathbf{S_1}$, …, $\mathbf{S_n}$ be $n$ random vectors of $q$ dimensions. Such collection defines the random variables for a sample of size $n$. Then, let $\mathbf{S_k} = (S_{k,1}, ..., S_{k,q})$ denotes the set of random variables describing the bio-tracer profile of the $k^{th}$ individual of the profile and $\mathbf{s_k} = (s_{k,1}, ..., s_{k,q})$ the observed values.

#### Bayesian approach

Let $[A_i|\mathbf{s}]$ be the probability that the individuals of the sample originate from the populations $i$ given the bio-tracers values observed. For any population $i$ ($i \in \mathbb{N}_p$):

$$[A_i|\mathbf{s}] = \frac{[A_i \cap \mathbf{s}]}{[s]}$$

Applying Bayes theorem yields:

$$[A_i|\mathbf{s}] = \frac{[\mathbf{s}|A_i]\,[A_i]}{[s]}$$

Assuming that $A_i, ..., A_p$ is a partition (i.e. those are the only potential population of origins):

$$[A_i|\mathbf{s}] = \frac{[\mathbf{s}|A_i]\,[A_i]}{\sum_j [\mathbf{s}|A_j]\,[A_j]} \tag{1}$$

using the bio-tracers distribution and assuming independence among individuals of the sample:

$$[\mathbf{s}|A_j] = \prod_k f_j(\mathbf{s_k}) \tag{2}$$

(1) becomes:

$$[A_i|\mathbf{s}] = \frac{\prod_k^n f_i(\mathbf{s_j})\,[A_i]}{\sum_j \prod_k f_j(\mathbf{s_k})\,[A_j]} \tag{3}$$

In case where bio-tracers are independent (3) can be written as:

$$f_j(\mathbf{s_k}) = \prod_l f_{j,l}(s_{k,l}) \tag{4}$$

which yields:

$$[A_i|\mathbf{s}] = \frac{[A_i]\prod_k \prod_l f_{i,l}(s_{k,l})}{\sum_j [A_j]\prod_k \prod_l f_{j,l}(s_{k,l})} \tag{5}$$

where the probability $[A_j]$ is a prior information. This can be used to complement the information about the sample. For instance, in seafood authentication, it can be used to reflect knowledge on species distribution or fisheries pressure. If no information is available, it is a reasonable assumption to consider populations equipossible in which case the relative size of population can be used as weights and in case population sizes are comparable (or no information is available) for any $(i, j)$, $[A_i] = [A_j]$, (5) then becomes:

$$[A_i|\mathbf{s}] = \frac{\prod_k \prod_l f_{i,l}(s_{k,l})}{\sum_j \prod_k \prod_l f_{j,l}(s_{k,l})} \tag{6}$$

Those assumptions are the same as the ones used by a Naive Bayesian Classifier (one of the three approaches we used in the main text).

### Effect of dissimilarity between 2 distributions

#### General considerations

Broadly speaking, there are three main directions that can lead to improving the inference process described above:

1. improving the *a priori* knowledge (which we do not consider here);
2. considering a larger quantity and/or higher quality of observations;
3. using more reliable inference techniques.

Below, we focus on how the data quantity and the properties of the distribution themselves influence the success of authentication. We do so as an attempt to better explain how increasing dimensionality can lead to better discriminatory power. To this end, we use simple cases where mathematical developments are fairly straightforwards.

We consider a simplified situation wherein one sample whose individuals come from one of two existing populations $A_1$ and $A_2$ are to be distinguished based on the distribution of one single bio-tracer. Under these assumptions, (5) becomes:

$$[A_1|\mathbf{s}] = \frac{[A_1]\prod_k^n f_{1,1}(s_{j,1})}{[A_1]\prod_k^n f_{1,1}(s_{j,1}) + [A_2]\prod_k^n f_{2,1}(s_{j,1})} \tag{7}$$

if we assume $[A_1] = [A_2]$ (*i.e.* a non-informative prior) and assuming $\prod_k^n f_{1,1}(s_{j,1}) > 0$, then (7) becomes: :

$$[A_1|\mathbf{s}] = \frac{1}{1 + \frac{\prod_k^n f_{2,1}(s_{j,1})}{\prod_k^n f_{1,1}(s_{j,1})}} \tag{8}$$

Assuming that the sample originates from $A_1$, then our goal is to understand how the dissimilarities between the two probability distributions impact the ratio $\frac{\prod_k^n f_{2,1}(s_{j,1})}{\prod_k^n f_{1,1}(s_{j,1})}$ and thus $[A_1|\mathbf{s}]$. As the latter probability also describes the success rate of which a sample is ascribed to its true provenance, we sometimes refer to this quantity as *performance*. Note that because the size of the sample $n$ strongly influences the performance, we also examine $n$ affect $[A_1|\mathbf{s}]$.

We further simplify the problem and posit that $f_{1,1}$ is the density function of a Normal distribution $\mathcal{N}(\mu_1, \sigma_1)$ and $f_{1,2}$ is the density function of Normal distribution $\mathcal{N}(\mu_2, \sigma_2)$. (7) becomes:

$$[A_1|\mathbf{s}] = \frac{1}{1 + \left(\frac{\sigma_1}{\sigma_2}\right)^n \exp\left(-\frac{1}{2}\left(\sum_{k=1}^n \left(\frac{s_k-\mu_2}{\sigma_2}\right)^2 - \left(\frac{s_k-\mu_1}{\sigma_1}\right)^2\right)\right)} \tag{9}$$

In the two following sections, we will consider two cases where dissimilarity is straightforwardly defined :

1. $\sigma_1 = \sigma_2 = \sigma$ where the dissimilarity will be quantified as $|\mu_1 - \mu_2|$;
2. $\mu_1 = \mu_2 = \mu$ where the dissimilarity will be quantified as $\frac{\sigma_1}{\sigma_2}$.

#### Identical variances

Under the assumption that $\sigma_1 = \sigma_2 = \sigma$, (9) becomes:

$$[A_1|\mathbf{s}] = \frac{1}{1 + \exp\left(-\frac{1}{2\sigma^2}\left(\sum_{k=1}^n (s_k - \mu_2)^2 - (s_k - \mu_1)^2\right)\right)} = \frac{1}{1 + \exp(-R)} \tag{10}$$

where:

$$R = \frac{1}{2\sigma^2}\sum_{k=1}^n (s_k - \mu_2)^2 - (s_k - \mu_1)^2 \tag{11}$$

Expanding $R$ we obtain:

$$R = \frac{1}{2\sigma^2}\sum_{k=1}^{n}(s_k - \mu_2 + s_k - \mu_1)(s_k - \mu_2 - s_k + \mu_1) = \frac{1}{2\sigma^2}\left(n(\mu_2^2 - \mu_1^2) + 2(-\mu_2 + \mu_1)\sum_{k=1}^{n} s_k\right)$$

We are now looking for an expression of the expected performance, i.e. $\mathbb{E}\left([A_1|\mathbf{s}]\right)$ for any $n$. As we assume that all individuals of the sample originate from $A_1$, if $X_n = \sum_{k=1}^{n} s_k$ then we have $X_n \sim \mathcal{N}(n\mu_1, \sqrt{n}\sigma)$. As $\mathbb{E}(X_n) = n\mu_1$ one can notice that

$$\mathbb{E}(R) = \left(n(\mu_2^2 - \mu_1^2) + 2n\mu_1(-\mu_2 + \mu_1)\right) = \frac{n}{2\sigma^2}(\mu_1 - \mu_2)^2$$

On a side note, as for all $x > 0$, $x \to \frac{x}{1+exp(-x)}$ is concave, following Jensen's inequality, we obtain the following upper boundary:

$$\mathbb{E}\left([A_1|\mathbf{s}]\right) \leq \frac{1}{1 + \exp\left(-\frac{n}{2}\left(\frac{\mu_1 - \mu_2}{\sigma}\right)^2\right)}$$

For the exact computation of $\mathbb{E}([A_1|\mathbf{s}])$, given that $X_n \sim \mathcal{N}\left(n\mu_1, \sqrt{n}\sigma\right)$, then $R \sim \mathcal{N}\left(\frac{n}{2\sigma^2}(\mu_2 - \mu_1)^2, \frac{\sqrt{n(\mu_1-\mu_2)^2}}{\sigma}\right)$. Using (10), we conclude that $[A_1|\mathbf{s}] \sim \mathit{Logitnormal}\left(\mu_{LN}, \sigma_{LN}\right)$ where $\mu_{LN} = \frac{n}{2\sigma^2}(\mu_2 - \mu_1)^2$ and $\sigma_{LN} = \frac{\sqrt{n(\mu_1-\mu_2)^2}}{\sigma}$. The moments do not have closed forms (Frederic and Lad 2008) but are straightforwardly computed numerically (see Wutzler 2018). Importantly enough here $\frac{\mu_{LN}}{\sigma_{LN}} = \frac{1}{2}\sigma_{LN}$, so the ratio increases with $\sigma_{LN}$ which is a monotonically increasing function of $|\mu_1 - \mu_2|$ and $n$. Following Frederic and Lad (2008), we therefore conclude that $\mathbb{E}([A_1|\mathbf{s}])$ increases with $|\mu_1 - \mu_2|$ and $n$ and illustrate this in Fig. S1.

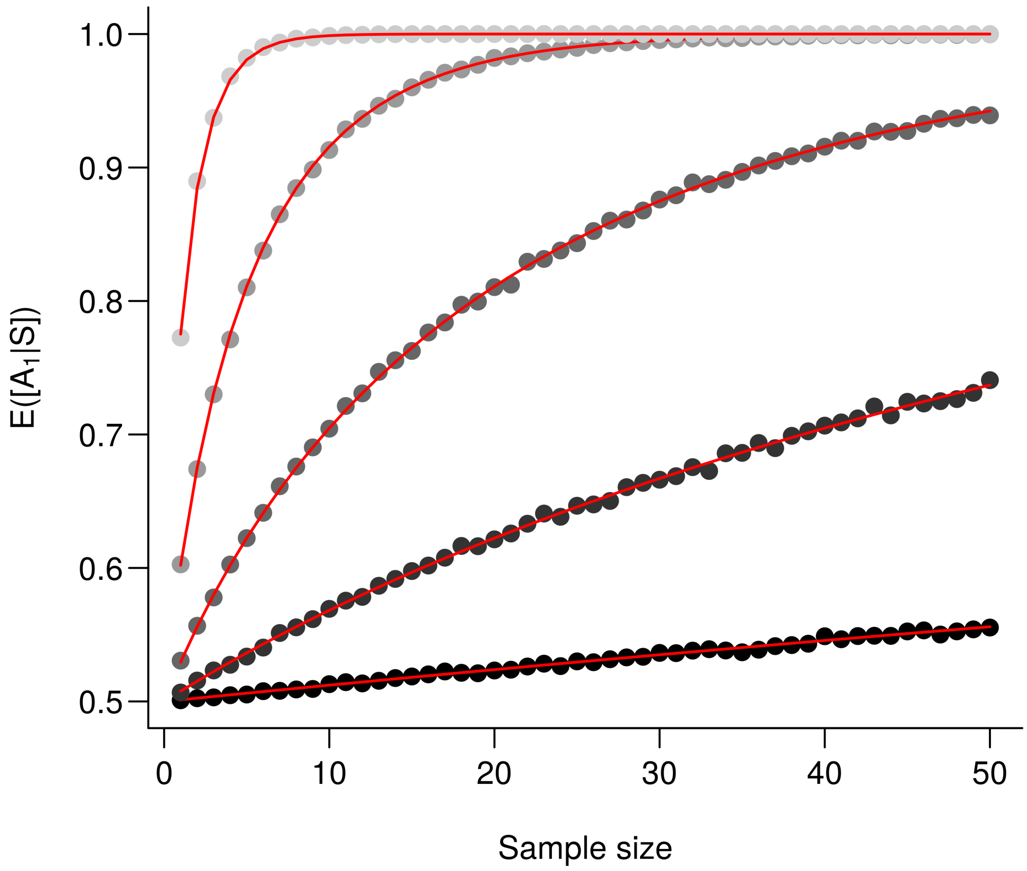

Fig. S1: **The larger the difference between the means the better the performance**. $\mathbb{E}\left([A_1|\mathbf{s}]\right)$ is computed for an increasing value $(\mu_1-\mu_2)^2$. Every point corresponds to the average value from 10,000 simulations and red lines correspond to the analytic solutions. From the darkest to the lightest gray, triplets $\{\mu_1,\mu_2,\sigma\}$ are as follows: $\{0,0.1,1\}$, $\{0,0.25,1\}$, $\{0,0.5,1\}$, $\{0,1,1\}$ and $\{0,1,0.5\}$.

#### Identical means

Now we examine $\mathbb{E}([A_1|\mathbf{s}])$ for $\mu_1=\mu_2=\mu$. Under such condition, (9) becomes:

$$[A_1|\mathbf{s}] = \frac{1}{1+\left(\frac{\sigma_1}{\sigma_2}\right)^n \exp\left(-\frac{1}{2}\left(\frac{1}{\sigma_2^2}-\frac{1}{\sigma_1^2}\right)\sum_{k=1}^n (s_k-\mu)^2\right)} \tag{12}$$

Assuming $X\sim\mathcal{N}(\mu,\sigma_1)$, by defining $Y=\sum_{k=1}^n\left(\frac{X-\mu}{\sigma_1}\right)^2$ we have $Y\sim\chi^2(n)$ and therefore:

$$\mathbb{E}\left([A_1|\mathbf{s}]\right) = \int_0^{+\infty} \frac{1}{1+\left(\frac{\sigma_1}{\sigma_2}\right)^n \exp\left(-\frac{1}{2}\left(\frac{\sigma_1^2}{\sigma_2^2}-1\right)y\right)} f(y,n)dy$$

where $f$ is the probability density function of $Y$. Using the expression of $f$, we obtain:

$$\mathbb{E}\left([A_1|\mathbf{s}]\right) = \int_0^{+\infty} \frac{\left(\frac{1}{2}\right)^{\frac{n}{2}} \frac{1}{\Gamma(n/2)} y^{\frac{n}{2}-1} \exp(\frac{-y}{2})}{1 + \left(\frac{\sigma_1^2}{\sigma_2^2}\right)^{\frac{n}{2}} \exp\left(-\frac{1}{2}\left(\frac{\sigma_1^2}{\sigma_2^2} - 1\right) y\right)} dy \tag{13}$$

where $\Gamma$ is the gamma function. At first glance, it is hard to determine how $\mathbb{E}([A_1|\mathbf{s}])$ changes with with $n$ and $\frac{\sigma_1}{\sigma 2}$. To examine this, we posit:

$$h(x, n) = \int_0^{+\infty} g(y, x, n) f(y, n) dy$$

where

$$g(y, x, n) = \frac{1}{1 + x^n \exp\left(-\frac{1}{2}(x^2 - 1)y\right)}$$

One can trivially shows that for any $n > 1$, if $x = 1$ then $h(x, n) = \frac{1}{2}$. We further conjecture that for any $x \in \mathbb{R}$, $x \to h(\exp(x), n)$ is symmetric and monotonically increasing on $x \in \mathbb{R}^+$ increase. We also conjecture that for any reals $n > 1$ and $m > 1$, if $m > n$ then $h(x, m) > h(x, n)$. Simulations presented in Fig. S2 (a) support our conjectures. Keeping this in mind, we computed $\mathbb{E}\left([A_1|\mathbf{s}]\right)$ for various values and an increasing sample size in Fig. S2 (b).

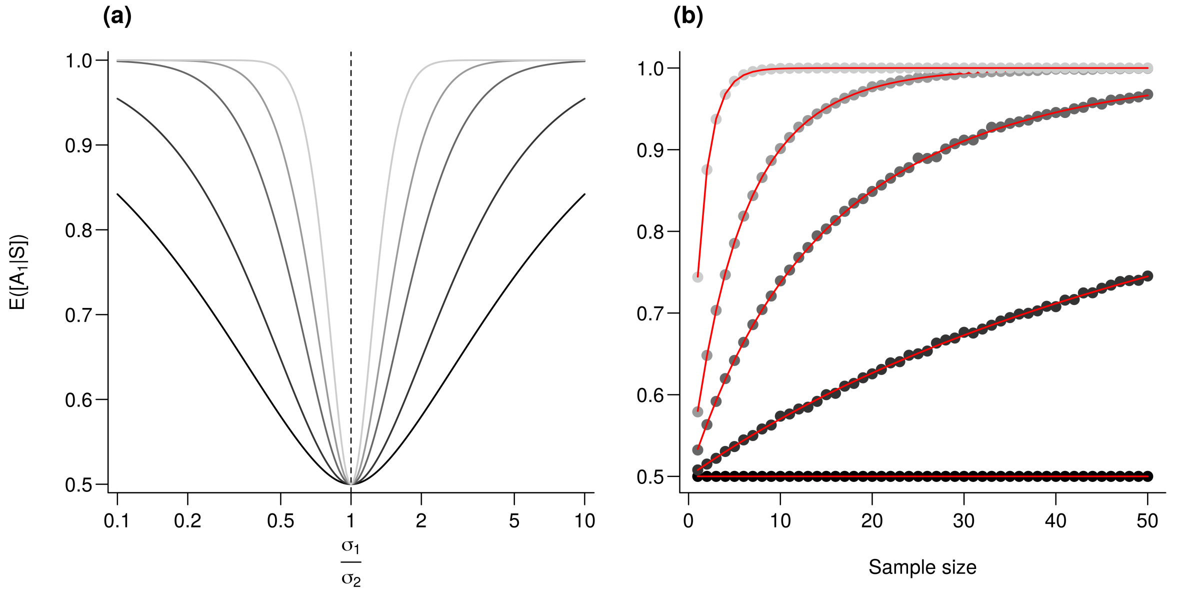

Fig. S2: **Effect of the variance ratio on performances**. **(a)**, $\mathbb{E}\left([A_1|\mathbf{s}]\right)$ is computed for an increasing $\frac{\sigma_1}{\sigma_2}$ value. From the darker to the lighter gray, $n$ (sample size) increases as follows : 1, 2.5, 5, 10, and 25. **(b)**), $\mathbb{E}\left([A_1|\mathbf{s}]\right)$ is computed for an decreasing ratio $\frac{\sigma_1}{\sigma_2}$. Every point represents the average value from 10,000 simulations, red lines correspond to the analytic solutions. For all simulations, $\mu_1 = \mu_2 = 0$, $\sigma_1 = 1$ and from the darkest to the lightest gray, $\sigma_2$ increases as follows : 1, 1.2, 1.5, 2 and 5.

### Effect of dimensionality

#### General considerations

In this section, we explore how increasing the number of bio-tracers $q$ affects the inference. As we previously did, we consider only two populations and we posit that the sample originates from $A_1$. Without further assumption on the co-distribution of bio-tracers, (8) becomes:

$$[A_1|\mathbf{s}] = \frac{1}{1 + \frac{\prod_k^n f_2(s_k)}{\prod_k^n f_1(s_k)}} \tag{14}$$

,

where $f_1$ and $f_2$ are $q$-variate density functions. We now have $q$ sets of random variables: $(S_{1,1}, ..., S_{1,n}), \ldots,$ $(S_{q,1}, ..., S_{q,n})$. Under the assumption of independence among individuals of the sample:

$$\forall (i,j) \in \mathbb{N}_n^2, i \neq j, [S_{1,i} \cap S_{1,j}] = [S_{1,i}][S_{1,j}]$$

Note that in the general case, adding bio-tracers improves the inference process only if

$$\prod_{k=1}^{n} \frac{f_{2,1}(s_k)}{f_{1,1}(s_k)} \geqslant \prod_{k=1}^{n} \frac{f_2(s_k)}{f_1(s_k)} \tag{15}$$

It is hard to assert whether it is generally true and below we use simple case to discuss under what conditions this holds true.

#### Simplification

We first assume that the bio-tracers are independent, in this case we have

$$\prod_{k=1}^{n} \frac{f_2(s_k)}{f_1(s_k)} = \prod_{k=1}^{n} \prod_{l=1}^{q} \frac{f_{2,l}(s_{j,l})}{f_{1,l}(s_{j,l})}$$

which is almost equivalent to increasing the number of samples and demonstration are similar to the previous ones. For instance, if for any $l$, $X_{n,l} \sim \mathcal{N}\left(n\mu_{1,l}, \sqrt{n}\sigma\right)$, then (11) becomes

$$R = \frac{1}{2\sigma^2} \sum_{l=1}^{q} \sum_{k=1}^{n} (s_{k,l} - \mu_{2,l})^2 - (s_{k,l} - \mu_{1,l})^2 \tag{16}$$

which yields:

$$R = \frac{1}{2\sigma^2} \sum_{l=1}^{q} \left( n(\mu_{2,l}^2 - \mu_{1,l}^2) + 2(-\mu_{2,l} + \mu_{1,l}) \sum_{k=1}^{n} s_{k,l} \right) \tag{17}$$

Assuming $X_{n,l} = \sum_{k=1}^{n} s_{k,l}$, then we have $X_{n,l} \sim \mathcal{N}(\mu_{1,l}, \sqrt{n}\sigma)$, for any $l$. As all $X_{n,l}$ are independent, then $R \sim \mathcal{N}\left( \frac{n}{2\sigma^2} \sum_{l=1}^{q} (\mu_{2,l} - \mu_{1,l})^2, \frac{\sqrt{n}}{\sigma} \sqrt{\sum_{l=1}^{q} \left(\mu_{2,l} - \mu_{1,l}\right)^2} \right)$, thus $[A_1|\mathbf{s}] \sim$ *Logitnormal* $(\mu_{LN}, \sigma_{LN})$ where $\mu_{LN} = \frac{n}{2\sigma^2} \sum_{l=1}^{q} (\mu_{2,l} - \mu_{1,l})^2$ and $\sigma = \frac{\sqrt{n}}{\sigma} \sqrt{\sum_{l=1}^{q} \left(\mu_{2,l} - \mu_{1,l}\right)^2}$. Just as in the previous section *Identical* *variances*, we have $\frac{\mu_{LN}}{\sigma_{LN}} = \frac{1}{2}\sigma_{LN}$, so the ratio increases with $\sigma_{LN}$ which is a monotonically increasing function of $\sum_{l=1}^{q}(\mu_{2,l} - \mu_{1,l})^2$ and $n$ and thus, following Frederic and Lad (2008), combining bio-tracer (i.e. augmenting the number of terms in the sum) increases the performances (see Fig. S3). Note that in the simplified situation where $(\mu_{2,l} - \mu_{1,l})^2 = d$ for any $l$, $\sum_{l=1}^{q}(\mu_{2,l} - \mu_{1,l})^2 = qd$, then $q$ and $n$ and $d$ plays similar roles.

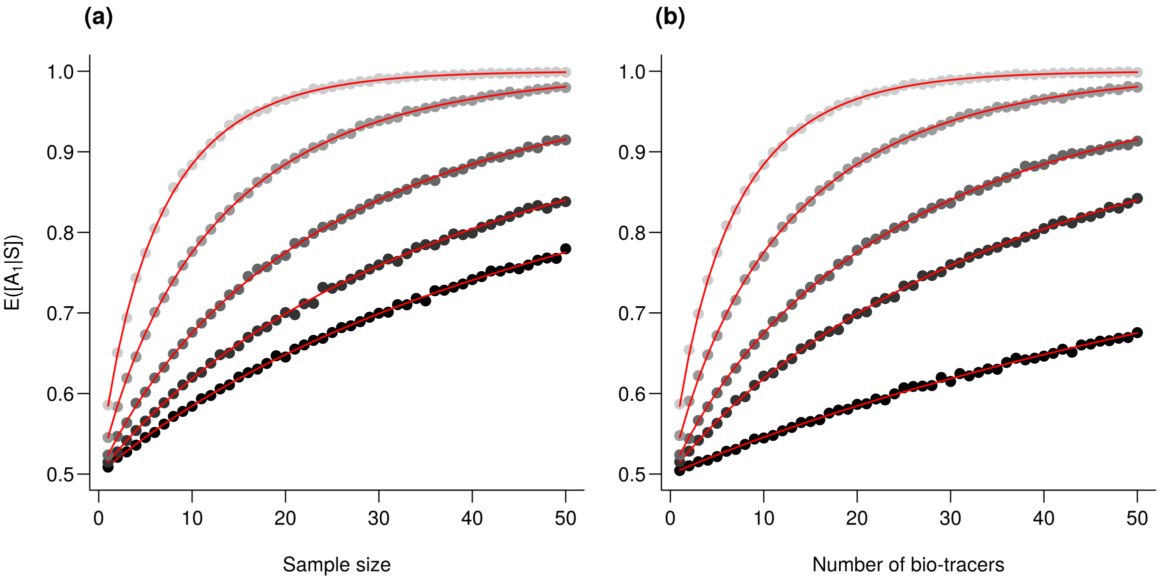

Fig. S3: **Effect of the number of bio-tracers combined on performances**. $\mathbb{E}([A_1|\mathbf{s}])$ is computed for an increasing number of samples **(a)** and for an increasing number of bio-tracers combined **(b)**. Every point represents the average value from 10,000 simulations, red lines correspond to the analytic solutions. For all simulations, $\sigma = 1$, for any real $l$, $\mu_{1,l} = 0$ and $\mu_{1,l} = 0.2$. **(a)**, from the darkest to the lightest gray, the number of bio-tracers combined are 2, 3, 5, 10 and 20, respectively. **(b)**, from the darkest to the lightest gray, the number of samples employed are 1, 3, 5, 10 and 20, respectively.

#### The role of correlation

When bio-tracers are not independent, we have the following relationship

$$\prod_{k=1}^{n}\frac{f_2(\mathbf{s_k})}{f_1(\mathbf{s_k})} = \prod_{k=1}^{n}\frac{f_{2,1}(s_{k,1})f_{2,S_{k,2}|S_{k,1}}(s_{k,2})...f_{2,S_{k,q}|S_{k,1},...,S_{k,q-1}}(s_{k,q})}{f_{1,1}(s_{k,1})f_{1,S_{k,2}|S_{k,1}}(s_{k,2})...f_{1,S_{k,q}|S_{k,1},...,S_{k,q-1}}(s_{k,q})}$$

where $f_{X|Y}$ denotes a conditional probability density function. We can only assert that as long as all ratios of conditional probability are less than 1, adding bio-tracers increase the overall performance, which should be true, on average.

That being mentioned, in the rest of the section, we examine the role of correlation among bio-tracers. We do so using a simple case where we assume that bio-tracer values are drawn in multivariate normal distributions, so for any population $j$, for any individual $i$, we have $\mathbf{S_{i,j}} \sim \mathcal{N}(\mu_\mathbf{j}, \mathbf{\Sigma_j})$, where $\mathbf{\Sigma_j}$ is the variance-covariance matrix.

We start by considering two bio-tracers. In this case, the general form of $\mathbf{\Sigma_j}$) is:

$$\mathbf{\Sigma_j} = \begin{bmatrix} \sigma_{1,j} & \text{cov}(S_{1,j}, S_{2,j}) \\ \text{cov}(S_{1,j}, S_{2,j}) & \sigma_{2,j} \end{bmatrix}$$

We assume that for any $i$ and $j$, $\sigma_{i,j} = 1$ and we focus on the role of $\text{cov}(S_{1,j}, S_{2,j})$. Note that under our assumption, $\text{cov}(S_{1,j}, S_{2,j})$ is also the correlation between the two bio-tracers ($\rho_j$). As we consider only two populations, we have two correlation values, $\rho_1$ and $\rho_2$, and we further simplify the problem by positing $\rho_1 = \rho_2$. In other words, we do not explore cases where different populations have different correlation structures among their bio-tracers. Therefore $\mathbf{\Sigma_1} = \mathbf{\Sigma_2} = \mathbf{\Sigma}$ that is of the following form:

$$\mathbf{\Sigma} = \begin{bmatrix} 1 & \rho \\ \rho & 1 \end{bmatrix}$$

To illustrate the role of $\rho$, we use (14) under the new set of assumptions and vary $\rho$ from 0.99 (bio-tracers are extremely correlated) to 0 (bio-tracers are independent) for an increasing sample size (Fig. S4). This shows that there is a continuum: from a situation where the two bio-tracers are very correlated and act effectively as a single bio-tracer, up to the situation where we have two uncorrelated bio-tracers, which performs better.

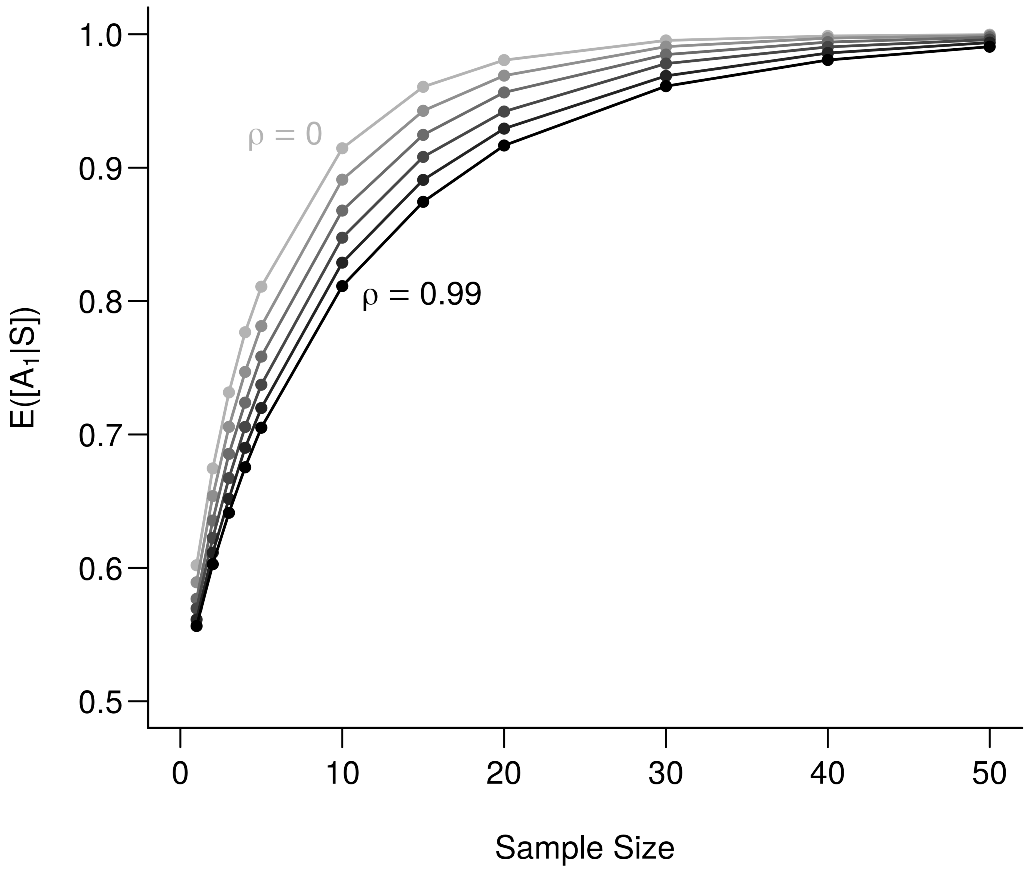

Fig. S4: **Effect of correlation among bio-tracers on the determination of origin**. The lighter the line the less correlated are the two bio-tracers. $\rho$ values range from 0 to 0.99 with an increment of 0.11, each point represents the average over 100,000 simulations.

In a second step and examine $[A_1|S]$ against an increasing dimensionality (*i.e.* a number of uncorrelated bio-tracers): from 1 to 10, for a sample of 25 individuals (Fig. S5). This shows that $\mathbb{E}([A_1|\mathbf{s}])$ increases with dimensionality.

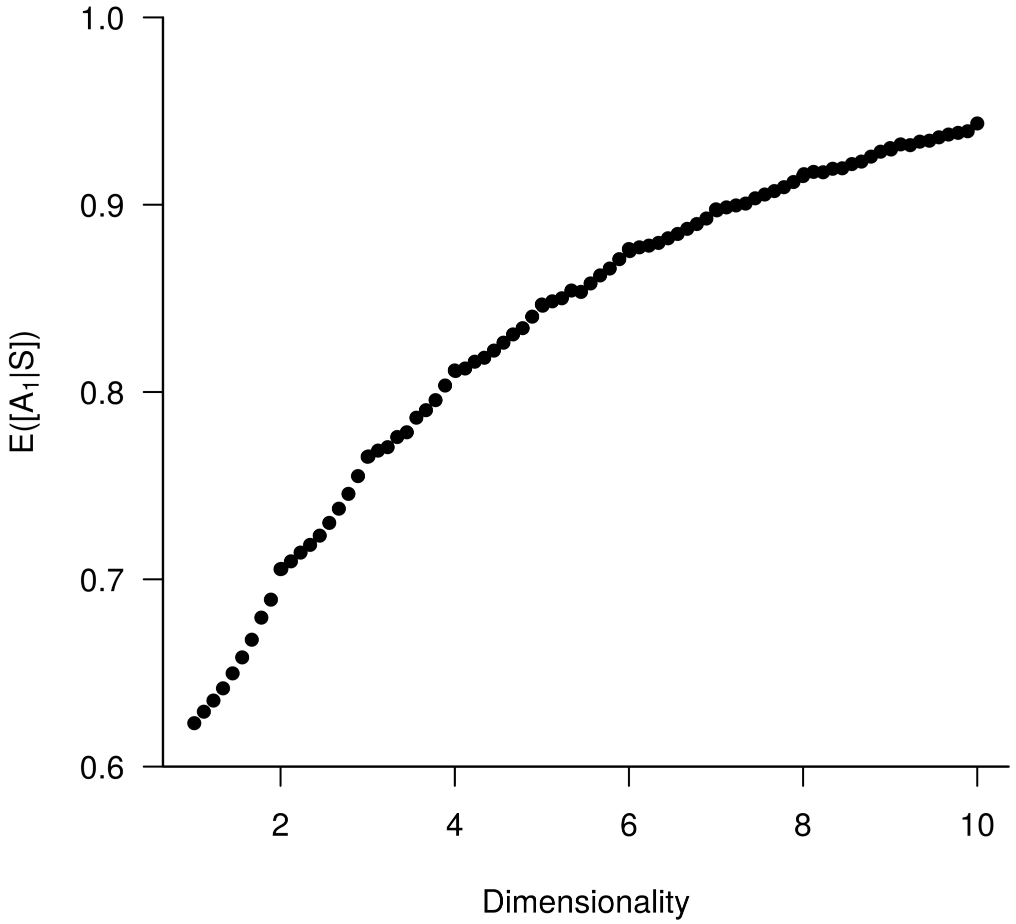

Fig. S5: **Effect of dimensionality of the set of bio-tracers on the determination of origin**. Each point represents the average value from 100,000 simulations.

### Conclusions

In order to strengthen the performance of the inference process herein described (i.e. in order to increase $\mathbb{E}([A_1|\mathbf{s}])$) and correctly determine the population of origin of one sample, one may:

1. increase the size of the sample,
2. use bio-tracers that maximize the dissimilarity of distributions,
3. increase the number of bio-tracers used (preferably as less correlated as possible),
4. use a combination of the factors aforementioned.

Note that in the above consideration we only consider two populations. When assuming more than two populations, mathematical developments are similar in principle but more tedious. For instance, if we extend the assumptions of section *Identical variances* to $m$ populations (normality and independence among the $l$ populations considered, $\sigma_l = 0$), 9 becomes:

$$[A_1|\mathbf{s}] = \frac{1}{1 + \sum_l \exp\left(-R_l\right)} \tag{18}$$

where $R_l \sim \mathcal{N}\left(\frac{n}{2\sigma^2}(\mu_l - \mu_1)^2, \frac{\sqrt{n(\mu_l - \mu_1)^2}}{\sigma}\right)$. Solving this analytically is beyond the scope of this appendix.

### References

Frederic, Patrizio, and Frank Lad. 2008. "Two Moments of the Logitnormal Distribution." *Communications in Statistics - Simulation and Computation* 37 (7): 1263–9. https://doi.org/10.1080/03610910801983178.

Wutzler, Thomas. 2018. *Logitnorm: Functions for the Logitnormal Distribution*.

### Supplementary Information

In this part of the Supplementary Information we added supplementary figures and supplemetary tables that complements the ones in the main text.

#### Influence of data augmentation and noise addition for Multi-Layer Perceptron (MLP)

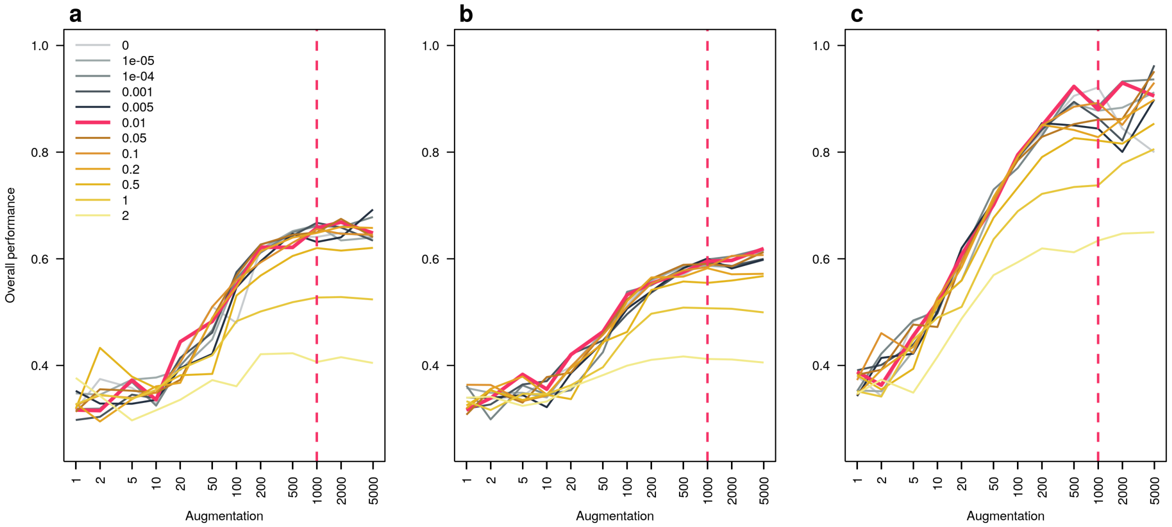

Fig. S6: **Effect of level augmentation and noise addition in the overall performance of the Multi-Layer Perceptron (MLP)**. Overall performance of the MLP are plotted against the number of times the data set is repeated (augmentation) and lines are colored according the noise level employed. We did so for the three stable isotopes we used **(a)**, the three first fatty acids **(b)** and all the 17 bio-tracers together **(c)**. The dotted red vertical line indicates the augmentation level we opted for and the red plain line represents the noise level we chose.

#### Influence of the training set size for the three methods

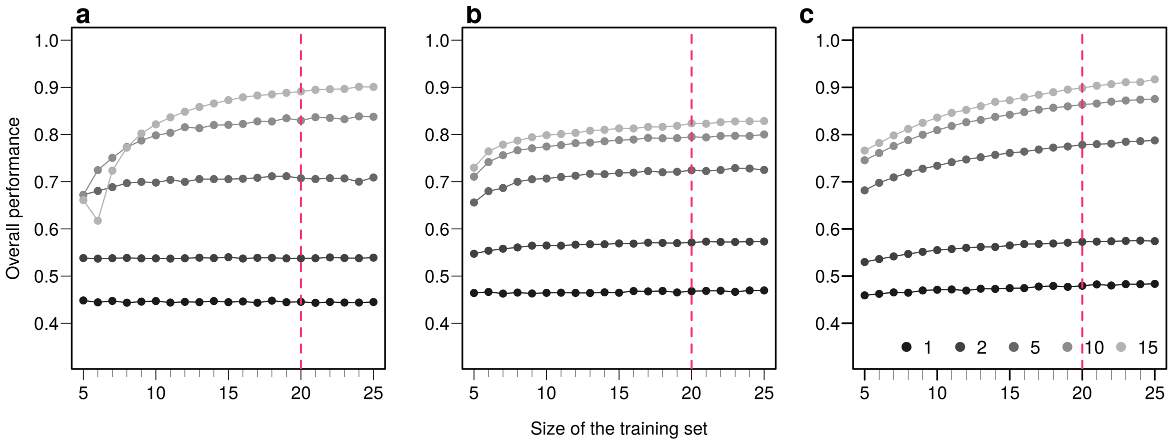

Fig. S7: **Effect of the size of the training set on overall performance of the three methods employed**. Overall performances are plotted for an increasing number of sample used in the training set. Points are colored according to the number of bio-tracers combined. Every point represents the average over up to 100,000 simulations (up to 500 axes combinations of bio-tracers and 200 replicates per combination). The vertical red dotted line indicates the size of the training set we used in the main text. The three panels correspond to three statistical approaches used: NBC (**a**), LDA (**b**), MLP (**c**). Note that potential gains in discriminatory power are increasing with the number of bio-tracers combined but obtaining such gains requires a larger number of samples.

### Results for pairs and triplets

#### Equivalent of figures 5-8 for the Naive Bayesian Classifier (NBC)

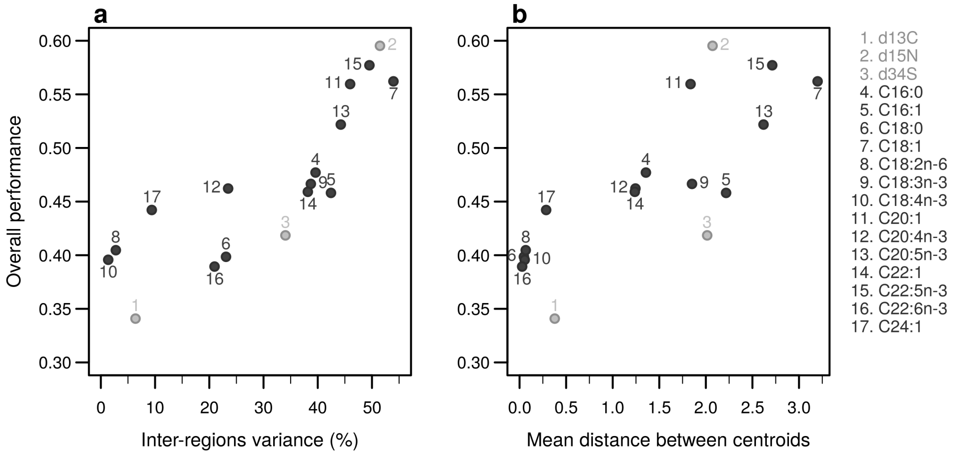

Fig. S8: **Performances of individual bio-tracers**. Overall performances of all 17 bio-tracers (listed on the right) using NBC are plotted against the proportion of inter-regions variance (**a**) and the mean distance between all pairs of region centroids (**b**).

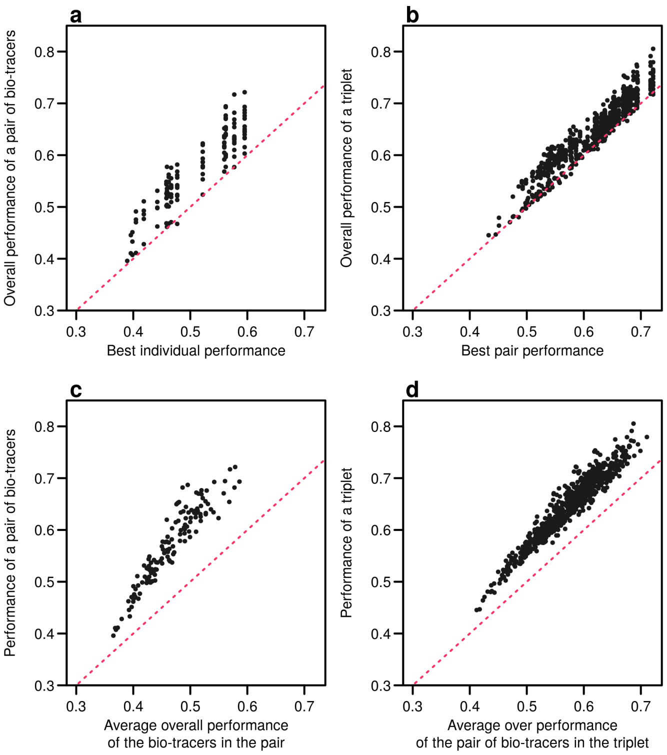

Fig. S9: **Including one more bio-tracers increases performance**. Overall performances of all pairs of bio-tracers using NBC are plotted against the best individual performing bio-tracers of the pair (**a**) and their average overall performance (**c**). Similarly, overall performances of all triplets of bio-tracers are plotted against the best performing pair of bio-tracers of the triplet (**b**) and their average overall performance. Magenta dashed lines represent the 1:1 slope.

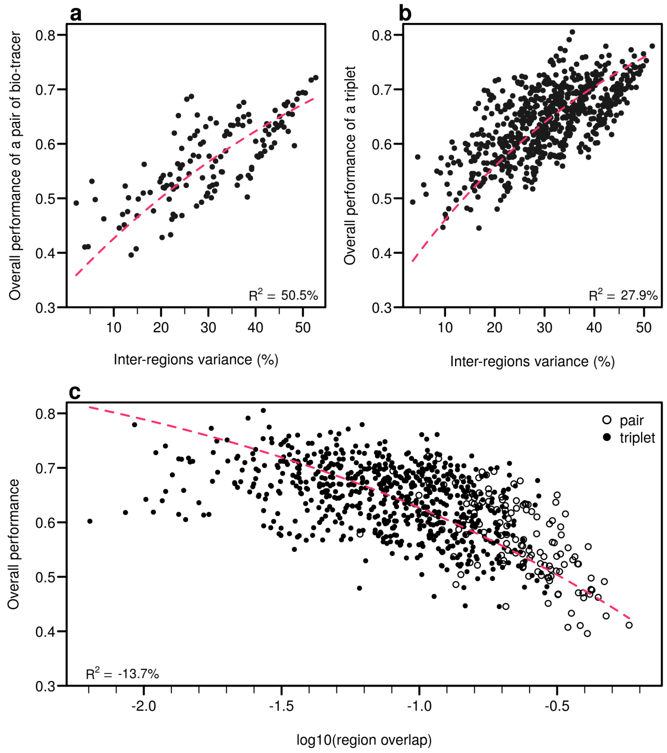

Fig. S10: **Efficient combinations of bio-tracers maximise inter-regional variance and minimize overlap of region data hypervolumes. a-b**, For all combinations of 2 (**a**) and 3 (**b**) bio-tracers, the overall performances (with NBC) of sets of bio-tracers are plotted against their inter-regions variance. **c**, We present relationship between the proportion of overlap of data between regions and the overall performances for all pairs and triplets of bio-tracers. Magenta dashed lines represent the results of non-linear leas-squares regression and the corresponding R-squared are added at the bottom of every panel.

#### Equivalent of figures 5-8 for Mutiple-Layer Perceptron (MLP, a class of Neural Network)

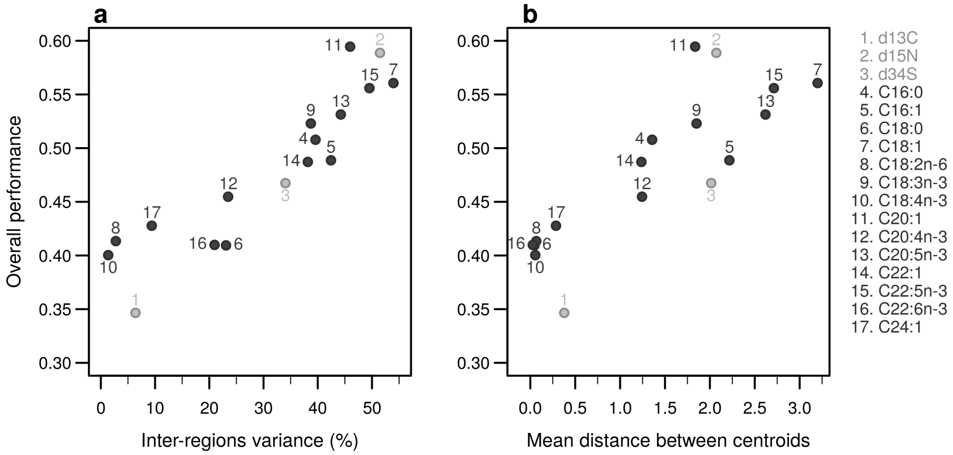

Fig. S11: **Performances of individual bio-tracers**. Overall performances of all 17 bio-tracers (listed on the right) using MLP are plotted against the proportion of inter-regions variance (**a**) and the mean distance between all pairs of region centroids (**b**).

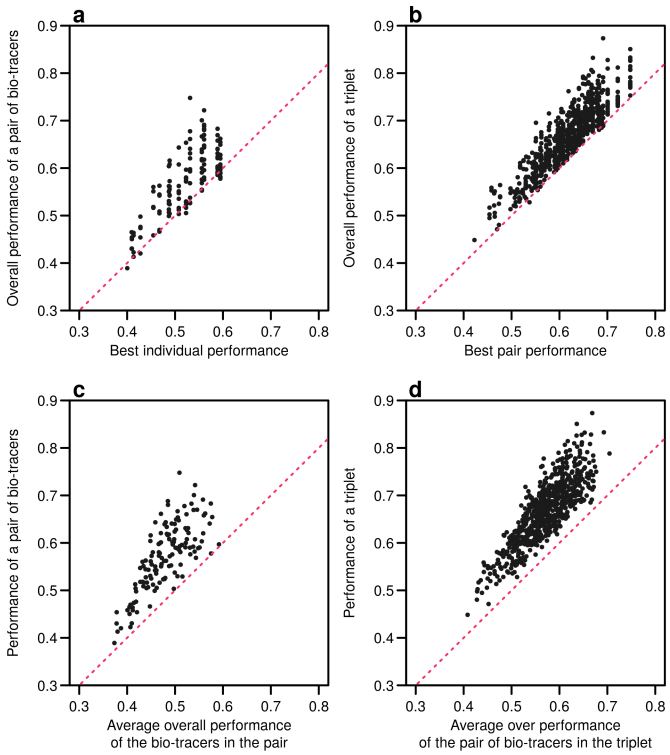

Fig. S12: **Including one more bio-tracers increases performance**. Overall performances of all pairs of bio-tracers using MLP are plotted against the best individual performing bio-tracers of the pair (**a**) and their average overall performance (**c**). Similarly, overall performances of all triplets of bio-tracers are plotted against the best performing pair of bio-tracers of the triplet (**b**) and their average overall performance. Magenta dashed lines represent the 1:1 slope.

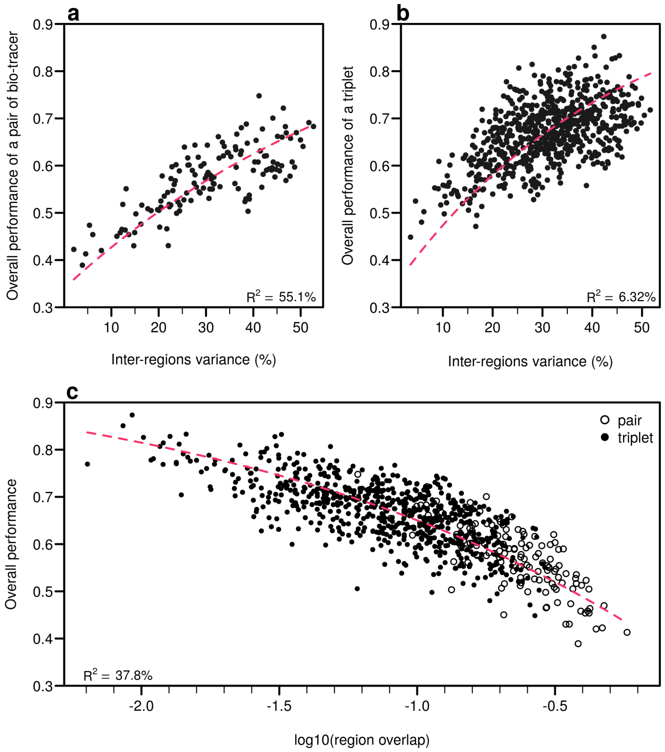

Fig. S13: **Efficient combinations of bio-tracers maximise inter-regional variance and minimize overlap of region data hypervolumes. a-b**, For all combinations of 2 (**a**) and 3 (**b**) bio-tracers, the overall performances (with MLP) of sets of bio-tracers are plotted against their inter-regions variance. **c**, We present relationship between the proportion of overlap of data between regions and the overall performances for all pairs and triplets of bio-tracers. Magenta dashed lines represent the results of non-linear leas-squares regression and the corresponding R-squared are added at the bottom of every panel.

Table S1: Top 10 pair of bio-tracers. Abbreviations are as follows: bt bio-tracer, op overall performance.

| Rank | LDA | | | NBC | | | MLP | | |
|---|---|---|---|---|---|---|---|---|---|
| | bt 1 | bt 2 | op | bt 1 | bt 2 | op | bt 1 | bt 2 | op |
| 1 | C20:5n-3 | C22:1 | 0.701 | $\delta^{15}N$ | C18:1 | 0.721 | C20:5n-3 | C22:1 | 0.748 |
| 2 | C18:1 | C22:6n-3 | 0.701 | C18:1 | C22:5n-3 | 0.717 | C18:1 | C18:3n-3 | 0.722 |
| 3 | C18:0 | C18:1 | 0.700 | C18:1 | C20:1 | 0.694 | C18:3n-3 | C22:5n-3 | 0.701 |
| 4 | C16:0 | C18:1 | 0.697 | $\delta^{15}N$ | C22:5n-3 | 0.693 | C18:1 | C22:5n-3 | 0.691 |
| 5 | $\delta^{15}N$ | C18:1 | 0.691 | C18:1 | C20:5n-3 | 0.693 | C18:0 | C18:1 | 0.688 |
| 6 | C20:1 | C22:6n-3 | 0.688 | $\delta^{15}N$ | C18:4n-3 | 0.687 | $\delta^{15}N$ | C18:1 | 0.683 |
| 7 | $\delta^{15}N$ | C16:1 | 0.687 | C18:4n-3 | C22:5n-3 | 0.682 | C16:0 | C18:1 | 0.683 |
| 8 | C18:1 | C20:1 | 0.677 | $\delta^{15}N$ | C20:1 | 0.682 | C18:1 | C22:6n-3 | 0.680 |
| 9 | C18:1 | C22:1 | 0.666 | C16:0 | C18:1 | 0.677 | C16:0 | C20:5n-3 | 0.678 |
| 10 | C18:1 | C20:5n-3 | 0.663 | $\delta^{15}N$ | C20:4n-3 | 0.676 | C18:1 | C20:5n-3 | 0.670 |

#### Best pairs and triplets of bio-tracers

Table S2: Top 10 triplet of bio-tracers. Abbreviations are as follows: bt bio-tracer, op overall performance.

| Rank | LDA | | | | NBC | | | | MLP | | | |
|---|---|---|---|---|---|---|---|---|---|---|---|---|
| | bt 1 | bt 2 | bt 3 | op | bt 1 | bt 2 | bt 3 | op | bt 1 | bt 2 | bt 3 | op |
| 1 | C18:1 | C20:5n-3 | C22:1 | 0.833 | $\delta^{15}N$ | C18:1 | C18:4n-3 | 0.805 | C18:1 | C20:4n-3 | C22:5n-3 | 0.873 |
| 2 | C18:1 | C20:5n-3 | C22:6n-3 | 0.793 | C18:1 | C18:4n-3 | C22:5n-3 | 0.791 | C18:3n-3 | C20:5n-3 | C22:1 | 0.851 |
| 3 | $\delta^{15}N$ | C20:5n-3 | C22:1 | 0.787 | $\delta^{15}N$ | C18:1 | C22:5n-3 | 0.779 | C18:1 | C20:5n-3 | C22:1 | 0.833 |
| 4 | C16:0 | C18:1 | C20:5n-3 | 0.771 | C18:1 | C20:4n-3 | C22:5n-3 | 0.779 | C18:1 | C20:4n-3 | C20:5n-3 | 0.832 |
| 5 | C18:0 | C18:1 | C20:4n-3 | 0.769 | C18:1 | C18:4n-3 | C20:5n-3 | 0.775 | $\delta^{15}N$ | C20:5n-3 | C22:1 | 0.828 |
| 6 | C18:1 | C20:4n-3 | C22:6n-3 | 0.769 | C18:1 | C18:3n-3 | C22:5n-3 | 0.773 | C18:3n-3 | C20:4n-3 | C22:5n-3 | 0.828 |
| 7 | C20:1 | C20:4n-3 | C22:6n-3 | 0.762 | $\delta^{15}N$ | C18:4n-3 | C22:5n-3 | 0.771 | C20:5n-3 | C22:1 | C22:5n-3 | 0.826 |
| 8 | C18:1 | C20:1 | C20:5n-3 | 0.759 | $\delta^{15}N$ | C18:1 | C20:5n-3 | 0.765 | C18:4n-3 | C20:5n-3 | C22:1 | 0.814 |
| 9 | C18:0 | C18:1 | C20:5n-3 | 0.758 | $\delta^{15}N$ | C18:1 | C20:4n-3 | 0.762 | C18:1 | C18:3n-3 | C20:4n-3 | 0.812 |
| 10 | $\delta^{15}N$ | C18:1 | C20:4n-3 | 0.755 | $\delta^{15}N$ | C18:1 | C18:3n-3 | 0.761 | C18:3n-3 | C22:5n-3 | C22:6n-3 | 0.810 |

#### DNA barcodes

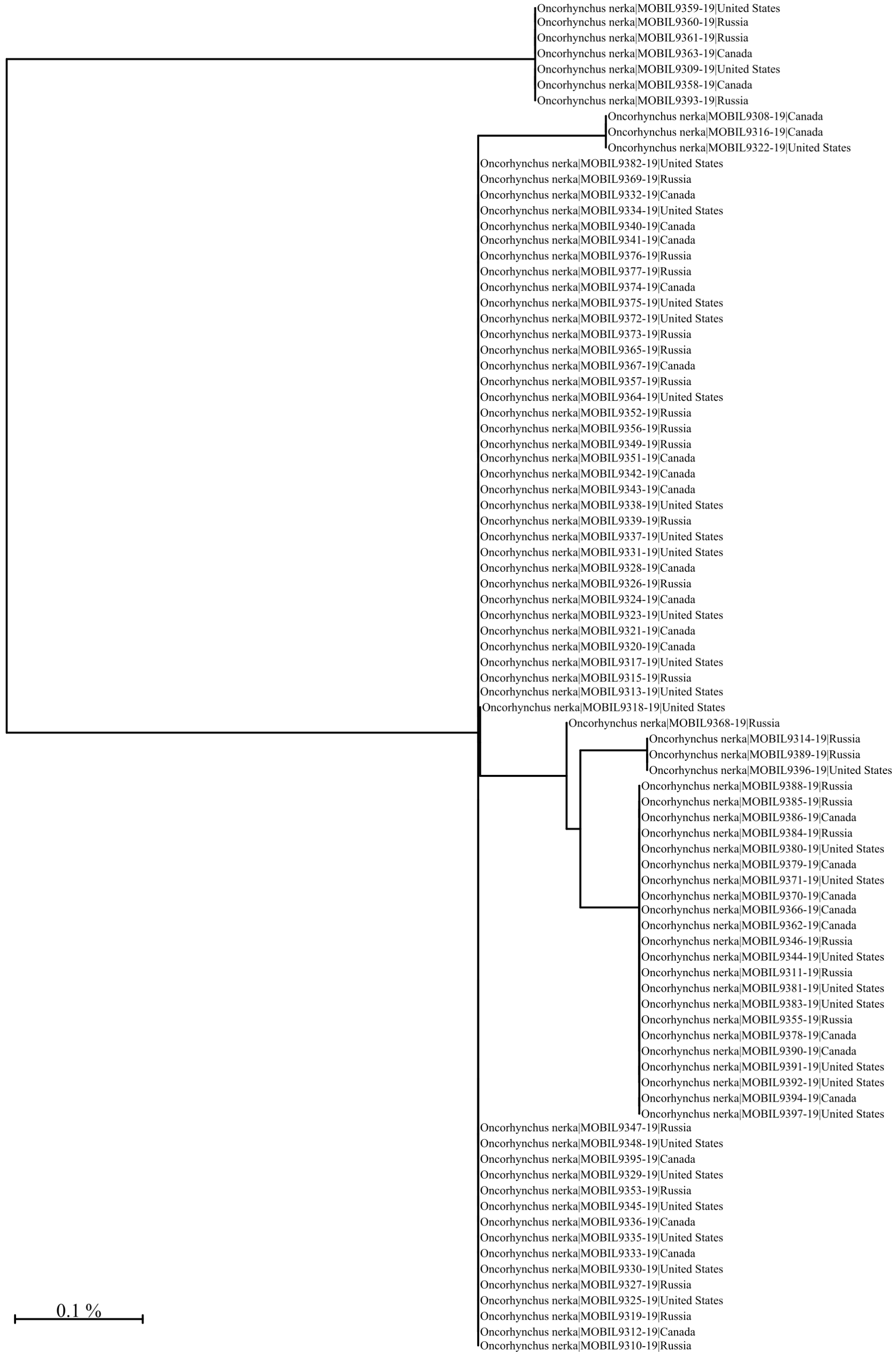

Fig. S14: Unrooted neighbour-joining tree (NJ) based on the p-distance of the 650 bp barcode region of the Cytochrome c Oxidase I gene. The NJ tree was generated using the Barcode of Life Data System V4 (Ratnasingham and Hebert 2007, see also boldsystems.org) using the Kimura 2 Parameter distance model (Kimura 1980) and sequences were generated using the LifeScanner DNA sequencing kit (lifescanner.net).

### References

Kimura, Motoo. 1980. "A Simple Method for Estimating Evolutionary Rates of Base Substitutions Through Comparative Studies of Nucleotide Sequences." *Journal of Molecular Evolution* 16 (2): 111–20. https://doi.org/10.1007/BF01731581.

Ratnasingham, Sujeevan, and Paul D. N. Hebert. 2007. "BARCODING: Bold: The Barcode of Life Data System (http://www.barcodinglife.org): BARCODING." *Molecular Ecology Notes* 7 (3): 355–64. https://doi.org/10.1111/j.1471-8286.2007.01678.x.
